# Stc1-expressing myofibroblasts are a developmentally distinct lineage cleared through intrinsic apoptosis in the neonatal lung

**DOI:** 10.1101/2025.06.12.659126

**Authors:** Melinda E. Snitow, Sylvia N. Michki, Fatima N. Chaudhry, Rachna Dherwani, Jeremy B. Katzen, David B. Frank, Jarod A. Zepp

## Abstract

Lung myofibroblasts are necessary for early postnatal alveolar growth and develop again during pathological fibrosis. Determining the unique contributions of multiple myofibroblast lineages to development and disease is hampered by a lack of genetic tools to distinguish between them. In this study, we generated a *Stc1^CreERT2^* mouse line that faithfully labels the developmentally transient secondary crest myofibroblasts (SCMF) and distinguishes SCMFs from alveolar duct myofibroblasts (DMF) and smooth muscle. SCMF populations expand by clonal proliferation of *Stc1*-expressing progenitors and contract by apoptosis. We deleted the intrinsic apoptosis effectors *Bax* and *Bak1* in the Stc1-lineage, which prevented SCMF clearance during alveologenesis. Single-cell RNA-seq revealed that residual Stc1-lineage cells lacking *Bax* and *Bak1* lose myofibroblast identity but express a combination of SCMF and DMF marker genes. Embryonic lineage tracing identified that SCMFs and DMFs have distinct progenitor populations with unique niches, and genetic activation of developmentally important signaling pathways could not interconvert these lineages. These findings establish Stc1-lineage SCMFs as a discrete population, developmentally divergent from DMFs, and define their life cycle in isolation from other myofibroblast lineages.

## INTRODUCTION

The transition to breathing air at birth is followed by a dramatic increase in the lung’s alveolar surface area, accomplished through the morphogenic process of postnatal alveologenesis. Alveolar sacs become encircled by transient myofibroblast rings that contract to promote secondary septa formation, thereby increasing the surface area for gas exchange. Understanding the origins and function of these secondary crest myofibroblasts (SCMF) could provide valuable insight into how myofibroblast identity is specified and possible cellular origins of lung disease (Zepp and Morrisey, 2019).

Single-cell RNA sequencing (scRNAseq) studies have shown that the lung mesenchyme comprises transcriptomically distinct cell types with unique anatomical localization. Recent work has highlighted that lung smooth muscle and myofibroblasts are arranged along the proximal-distal axis from the airway tree, extending outward toward the alveoli. Alveolar ducts are distal to the bronchioalveolar duct junction and are surrounded by myofibroblasts that span thick elastin fibers rather than smooth muscle (Chaudhry et al., 2024; Narvaez Del Pilar et al., 2022). SCMFs exist transiently in the distal alveolar regions, and their presence and ability to contract are required for alveolar maturation (Li et al., 2018; Li et al., 2020). SCMFs are depleted by the end of alveologenesis via apoptosis during alveolar wall thinning (Branchfield et al., 2016; Bruce et al., 1999; Schittny et al., 1998), and the remaining alveolar myofibroblast population is the recently recognized alveolar duct myofibroblast (DMF) that has a transcriptional signature distinct from SCMFs (Chaudhry *et al*., 2024; Narvaez Del Pilar *et al*., 2022). Prior studies on alveolar myofibroblasts could not distinguish between DMFs and SCMFs because the marker genes used (*Acta2*, *Pdgfra*, *Gli1*, *Fgf18*) label both populations (Chaudhry *et al*., 2024; Endale et al., 2017; Hagan et al., 2020; Li et al., 2015; Li *et al*., 2018; Moiseenko et al., 2017; Ntokou et al., 2015; Zepp et al., 2021). Without specific lineage tools, studies confounding DMFs with SCMFs leave open questions about their developmental relationship, unique biology, and relative contribution to pathological processes.

In this study, we generated a lineage tracing tool that distinguishes SCMFs from all other lung myofibroblasts, including DMFs. We analyzed the life cycle of the transient SCMF population, identified distinct embryonic progenitors of SCMF and DMF populations, and genetically deleted effectors of apoptosis, a key biological feature of SCMFs. We found that SCMF fate could not be shifted into the DMF lineage by blocking apoptosis or stimulating developmental signaling pathways key for myofibroblast development. These findings demonstrate that the Stc1-lineage+ SCMF lineage is distinct from DMFs, and highlight the specific biology of SCMFs.

## RESULTS

### *Stc1* is specific marker of SCMF in neonatal lung mesenchyme

To find a specific marker of SCMFs that distinguishes them from other myofibroblasts, we integrated three scRNA-seq datasets that sequenced mesenchyme at 12 developmental timepoints spanning SCMF development (Narvaez Del Pilar *et al*., 2022; Negretti et al., 2021; Zepp *et al*., 2021) (Figure 1A). Myofibroblast populations are transcriptomically similar and include airway smooth muscle (ASM), ductal myofibroblasts (DMF), and SCMFs. We examined these populations for novel marker genes specific to SCMFs and found that *Stc1* expression distinguishes SCMFs from other myofibroblasts (Figure 1B).

**Figure 1.**
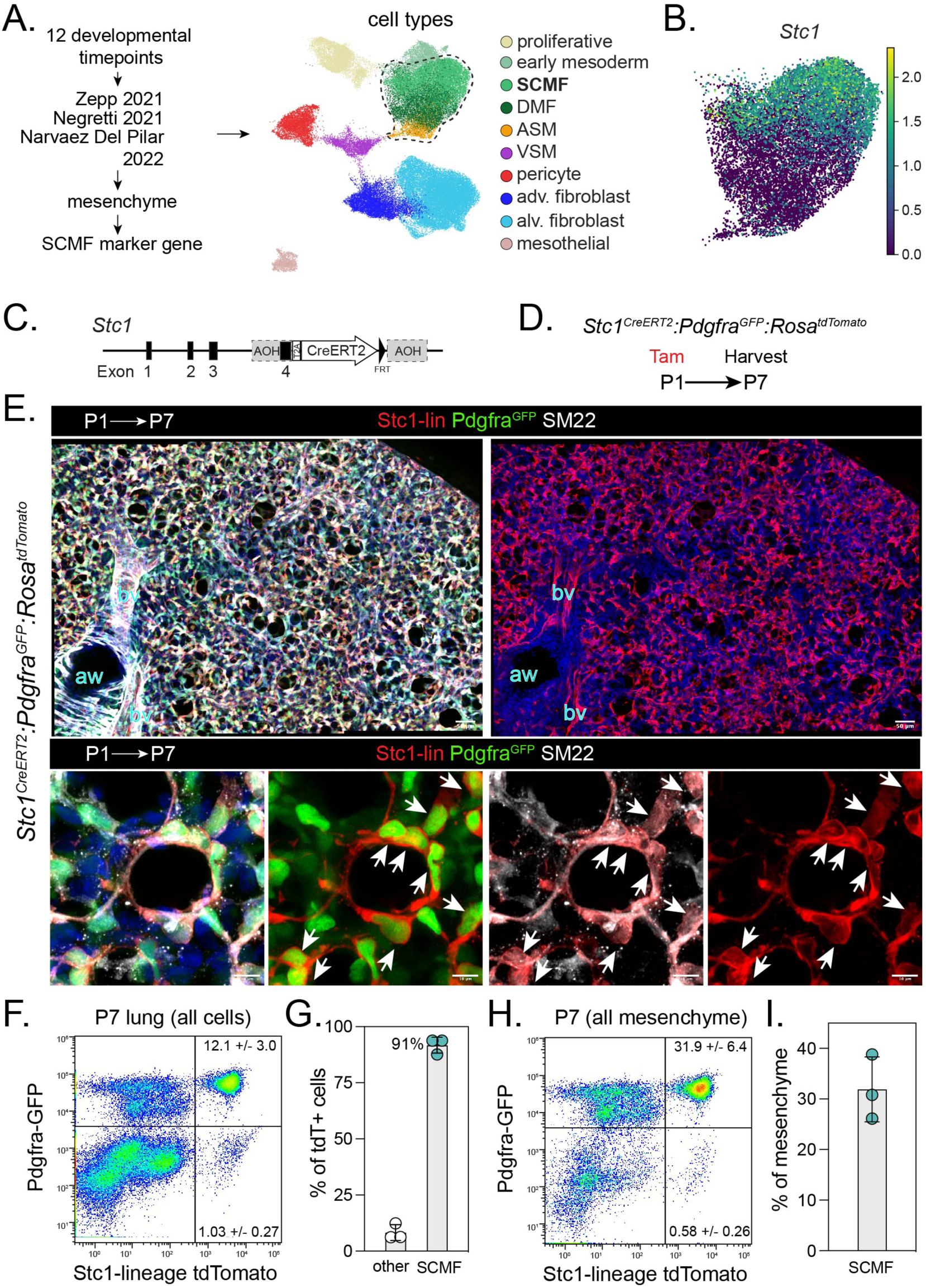
*Stc1* is a specific marker of SCMF in P7 lung mesenchyme. A) UMAP of lung mesenchyme at 12 developmental time points from 3 merged scRNA-seq datasets. Dashed line surrounds myofibroblast cell types. B) Gene expression of Stc1 in myofibroblast cell types from analysis in (A) at P3, P5, and P7. C) Diagram of *Stc1* locus targeted to generate a knock-in *Stc1^CreERT2^* allele. D) Experiment schematic for (E) through (I). E) Stc1-lineage labeled lung harvested at P7, and immunostained for SM22. Arrows indicate Stc1-lineage labeled cells that are also positive for Pdgfra^GFP^ and SM22. Z-projections of approximately 100 µm (upper) and 40 µm (lower). F-G) Flow cytometry analysis (F) and quantification (G) of all live cells in the P7 lung for Stc1-lineage tdTomato cells positive for Pdgfra^GFP^ (“SCMF”) or negative for Pdgfra^GFP^ (“other”). N = 3, mean ± SD. H-I) Flow cytometry analysis (H) and quantification (I) of mesenchymal cells in the P7 lung for Stc1-lineage tdTomato and Pdgfra^GFP^ (“SCMF”). N = 3, mean ± SD. Abbreviations: Stc1-lin, Stc1-lineage; aw, airway; bv, blood vessel; SCMF, secondary crest myofibroblast; DMF, ductal myofibroblast; ASM, airway smooth muscle; VSM, vascular smooth muscle; adv., adventitial; alv., alveolar. Scale bars: (E, top row) 50 µm, (E, bottom row) 10 µm.

We designed a knock-in *Stc1* lineage reporter by inserting a *CreERT2* sequence at the end of the last *Stc1* exon, separated by a T2A self-cleaving peptide sequence (Figure 1C, Supplementary Figure S1A,B). *Stc1-CreERT2* mice were crossed with a *R26R^LSL-tdTomato^* reporter for lineage tracing, and a *Pdgfra^GFP^*reporter to assess *Pdgfra* expression (Hamilton et al., 2003; Madisen et al., 2010). Neonates were dosed with tamoxifen on postnatal day 1 (P1) and harvested at P7 when SCMFs are abundant in the murine lung (Zepp *et al*., 2021). Mice untreated with tamoxifen showed no reporter expression (Supplementary Figure S1C). Stc1-lineage traced cells were abundant in the P7 lung, forming ring-like structures in the alveolar space and co-expressing SCMF markers SM22 and Pdgfra^GFP^ (Figure 1E).

We quantified the *Stc1* lineage by flow cytometry, finding that SCMFs double-positive for Stc1-lineage/Pdgfra^GFP^ comprise 12.1 ± 3.0% of all live cells in the P7 lung (Figure 1F, G). The remaining Stc1-lineage expressing cells negative for Pdgfra^GFP^ represented 1.03 ± 0.27% of all live cells, indicating that a minority of Stc1-lineage cells are non-SCMF and included arterial endothelium (Supplemental Figure S1D). Flow cytometry indicated that SCMFs account for 31.9 ± 6.4% of the lung mesenchyme (CD45/CD31/EpCAM-negative) at P7, with only 0.58 ± 0.26% of mesenchymal cells being labeled by Stc1-lineage alone, which included pericytes (Figure 1H, I and Supplementary Figure S1E, F).

### SCMFs expand clonally from Stc1-lineage progenitors and population contracts through apoptosis

SCMFs are transient myofibroblasts that have been observed to be largely depleted by the end of alveologenesis (Hagan *et al*., 2020; Li *et al*., 2018; Zepp *et al*., 2021). To better define the kinetics of SCMF development, we first assessed Stc1-lineage abundance at multiple timepoints during postnatal lung development between peak alveologenesis (P7) and post-alveologenesis (P30). This time course showed that the Stc1-lineage declines after P10 and is depleted by P30 (Figure 2A, B). Stc1-lineage cells had a corresponding increase in apoptosis as marked by cleaved Caspase 3 abundance between P10 and P15 (Figure 2C, D).

**Figure 2.**
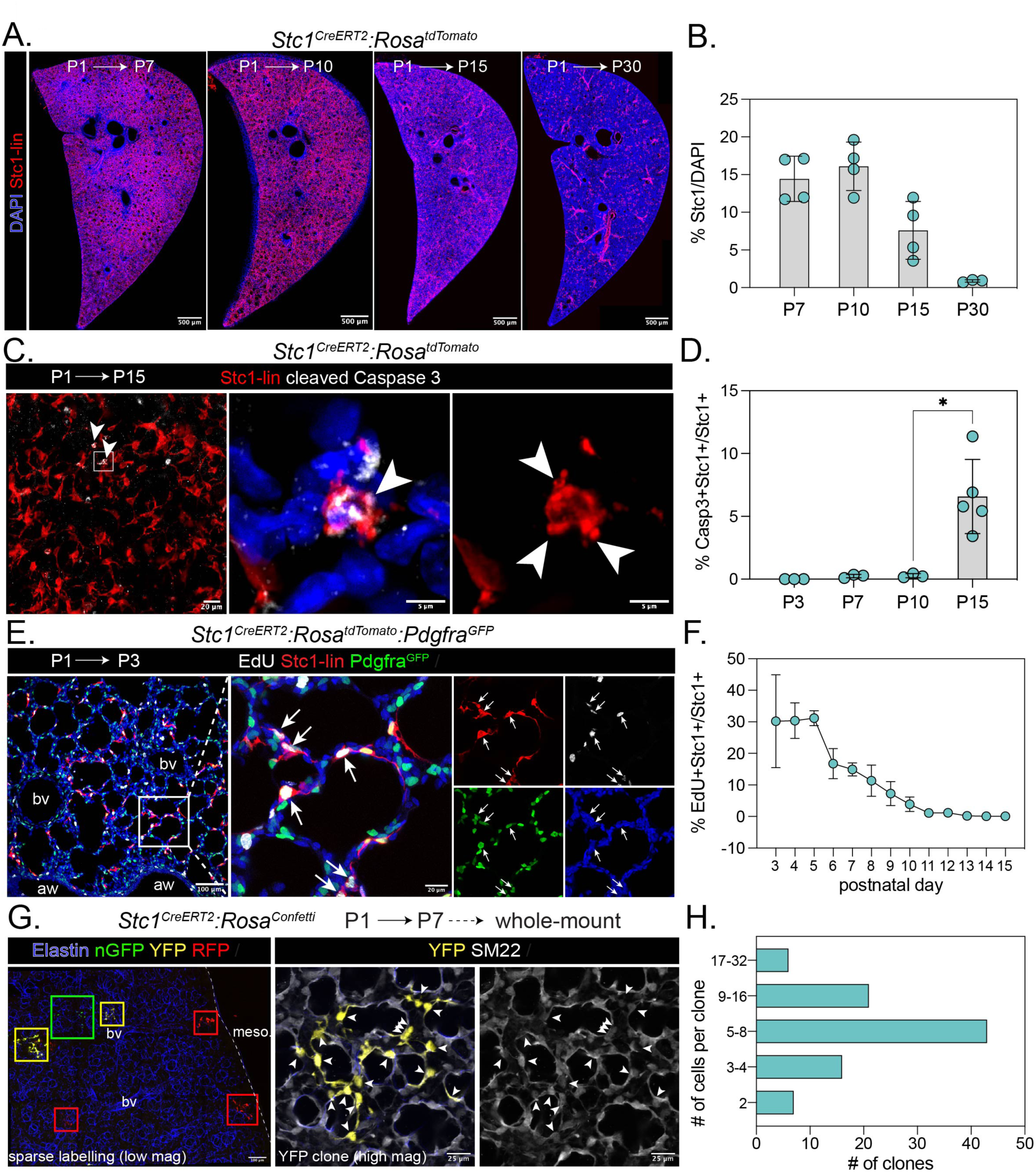
Stc1-lineage trace models SCMF depletion by apoptosis and SCMF expansion by progenitor division. A) Stc1-lineage trace from P1 through indicated days in left lung cross-sections. Z-projections of approximately 80 µm. B) Quantification of (A). C) Stc1-lineage trace from P1 through P15, co-immunostained with the apoptosis marker cleaved Caspase 3. Arrows in left and middle panels indicate Stc1-lineage cells that express cleaved Caspase 3; arrows in right panel indicate apoptotic bodies. Z-projections of approximately 130 µm. D) Quantification of (C). E) Stc1-lineage trace from P1 to P3, Pdgfra^GFP^, and EdU incorporation at P3. Arrows indicate triple-positive cells. Z-projections of approximately 10 µm (left) and 20 µm (inset). F) Quantification of (E). G) Stc1-lineage trace from P1 to P7 using the Rosa-confetti reporter in whole-mount accessory lobes. Colored rectangles indicate clones. Higher magnification of a YFP clone co-immunostained with SM22. Arrows indicate double-positive cells. Z-projections of approximately 250 µm (low mag) and 45 µm (high mag). H) Quantification of (G), using a histogram binned by log2. Abbreviations: Stc1-lin, Stc1-lineage; aw, airway; bv, blood vessel; meso, mesothelium; mag, magnification. Data are represented as mean and standard deviation in (B, D, F). (H) represents an absolute count of clones. Statistics: * *p* < 0.05. Scale bars: (A) 500 µm; (C, left) 20 µm; (C, middle and right) 5 µm; (E, left) 100 µm; (E, inset) 20µm; (G, left) 100 µm; (G, inset) 25 µm.

To characterize the dynamics of SCMF expansion between birth and peak abundance, we analyzed the proliferation of Stc1-lineage/Pdgfra^GFP^ cells by 5-ethynyl-2′-deoxyuridine (EdU) incorporation. EdU incorporation (4-hour pulse-chase) was highest in the earliest timepoints analyzed (P3-P5) during the period of low SCMF abundance, then declined starting at P6 as SCMF abundance peaked (Figure 2E, F). Proliferation reached its lowest point by P11, corresponding to the trend of reduced SCMF abundance and increased apoptosis. Postnatal proliferation dynamics differ between SCMFs, other Pdgfra^GFP^+ fibroblasts, and all lung cells, with proliferation peaking in these respective populations at P3-P5, P4, and P5 (Supplementary Figure S2A-D).

To characterize the progenitor capacity of individual SCMFs early in development, we performed clonal analysis by crossing *Stc1^CreERT2^* with the *Rosa^Confetti^*reporter (Snippert et al., 2010b). After tamoxifen administration at P1, we used whole-mount imaging of the intact accessory lobe to confirm sparse labeling of Stc1-lineage cells at P7 as indicated by spatially discrete single-color cell clusters that represent likely single-cell-derived clones (Figure 2G). We quantified the number of cells per clone from multiple litters and found that the majority of SCMF clones comprised 5-8 daughter cells (Stc1-lineage/SM22), indicating that progenitors underwent 3-4 cell divisions from P1 to P7 (Figure 2H). Together, these data characterize the proliferative expansion and apoptotic contraction of the SCMF population.

### Stc1-lineage excludes alveolar duct myofibroblasts

ASM, DMF, and SCMF are transcriptionally distinct myofibroblasts that occupy adjacent but physically discrete niches spanning the proximal to distal air space, respectively (Figure 3A, Supplementary Figure 3). We analyzed P7 SCMF and DMF marker genes from the aggregated scRNAseq atlases, finding that *Stc1* and *Hhip* expression can distinguish between these populations with few overlapping cells (Figure 3B).

**Figure 3.**
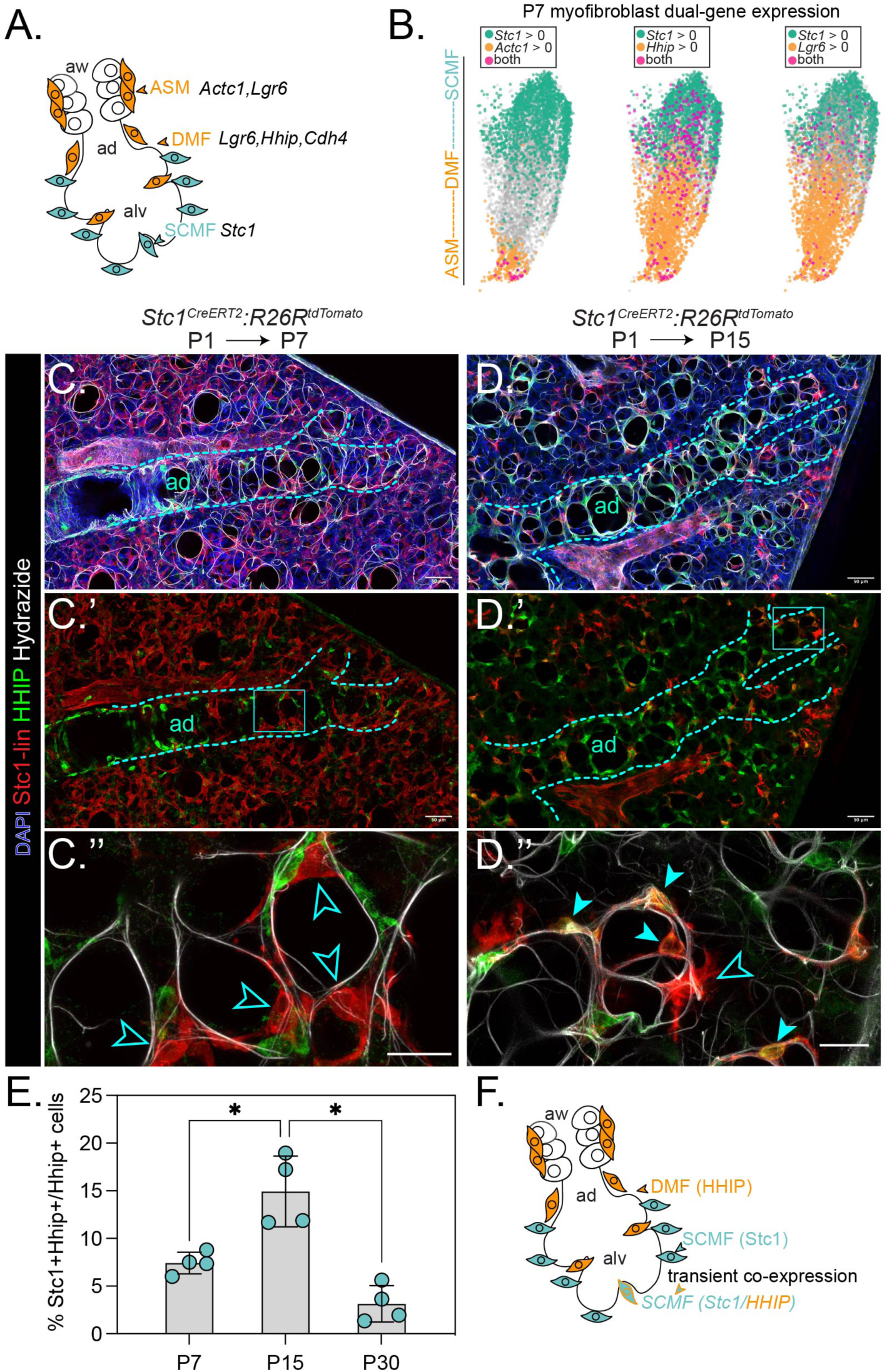
*Stc1*-lineage distinguishes SCMFs from ductal myofibroblasts. A) Diagram of myofibroblast cell types and marker genes surrounding the proximal to distal air spaces during alveologenesis. B) UMAP of scRNA-seq showing the myofibroblast populations, comparing cells expressing *Stc1* with cells expressing marker genes of the ASM and DMF populations shown in (A) using dual-gene expression (expression > 0 for indicated genes). C-D) Stc1-lineage trace from P1 to P7 (C) and P15 (D), co-immunostained with Hhip and hydrazide. Dashed lines outline alveolar ducts, rectangles (C’, D’) outline high magnification regions (C’’, D’’). Outlined arrows indicate Stc1-lineage traced cells that are negative for Hhip; filled arrows indicate double-positive cells. Z-projections of approximately 100 µm (C), 25 µm (C’’), and 40 µm (D, D’’). E) Quantification of cells Stc1-lineage traced at P1 that are positive for Hhip at the indicated ages. F) Model of distinct SCMF and DMF lineages, with some transient Hhip expression in some Stc1-lineage SCMFs. Abbreviations: Stc1-lin, Stc1-lineage; aw, airway; ad, alveolar duct; alv, alveolus; ASM, airway smooth muscle; DMF, ductal myofibroblast; SCMF, secondary crest myofibroblast. Scale bars: (C, C’, D, D’) 50 µm; (C’’, D’’) 20 µm. Graphs represent mean and standard deviation. Statistics: Kruskal-Wallis test statistic = 9.846, *p* = 0.0002. Post-hoc Mann-Whitney tests: *\* p <* 0.025.

To confirm that the Stc1-lineage distinguishes SCMFs from DMFs, we immunostained these tissues for the ASM/DMF marker HHIP at P7 and P15. While SCMFs were abundant at P7, they were distinct from the HHIP+ DMFs (Figure 3C). Interestingly, few co-expressing cells were transiently increased at P15 (Figure 3D-E). These cells tended to be at the distal end of alveolar ducts and resembled SCMF morphology, forming a ring-like structure around a thinner elastin ring (Figure 3D’, D’’). By P30, these HHIP/Stc1-lineage positive cells were depleted, suggesting that most double-positive cells were SCMFs that transiently expressed HHIP before apoptosis (Figure 3E, F). These lineage tracing results confirm that the *Stc1^CreERT2^* line effectively distinguishes myofibroblast sub-types along the proximal-distal axis.

### A conditional genetic knockout model of apoptosis prevents SCMF depletion

Next, we tested whether intrinsic apoptosis is required to deplete the SCMF population and to uncover the fate of SCMFs that are experimentally retained post-alveologenesis. We conditionally deleted the intrinsic apoptosis effectors, *Bax* and *Bak1*, in the Stc1-lineage, hereafter referred to as Stc1-BB Het and KO (Takeuchi et al., 2005) (Figure 4A). Between P16 and P30, Stc1-lineage+ cells are depleted in Stc1-BB Het controls, but were retained in Stc1-BB KO (Figure 4B, C). The retained Stc1-lineage cells were found throughout the distal lung, and most were morphologically compact, in contrast to their elongated appearance at P7 (Figure 4D-E).

**Figure 4.**
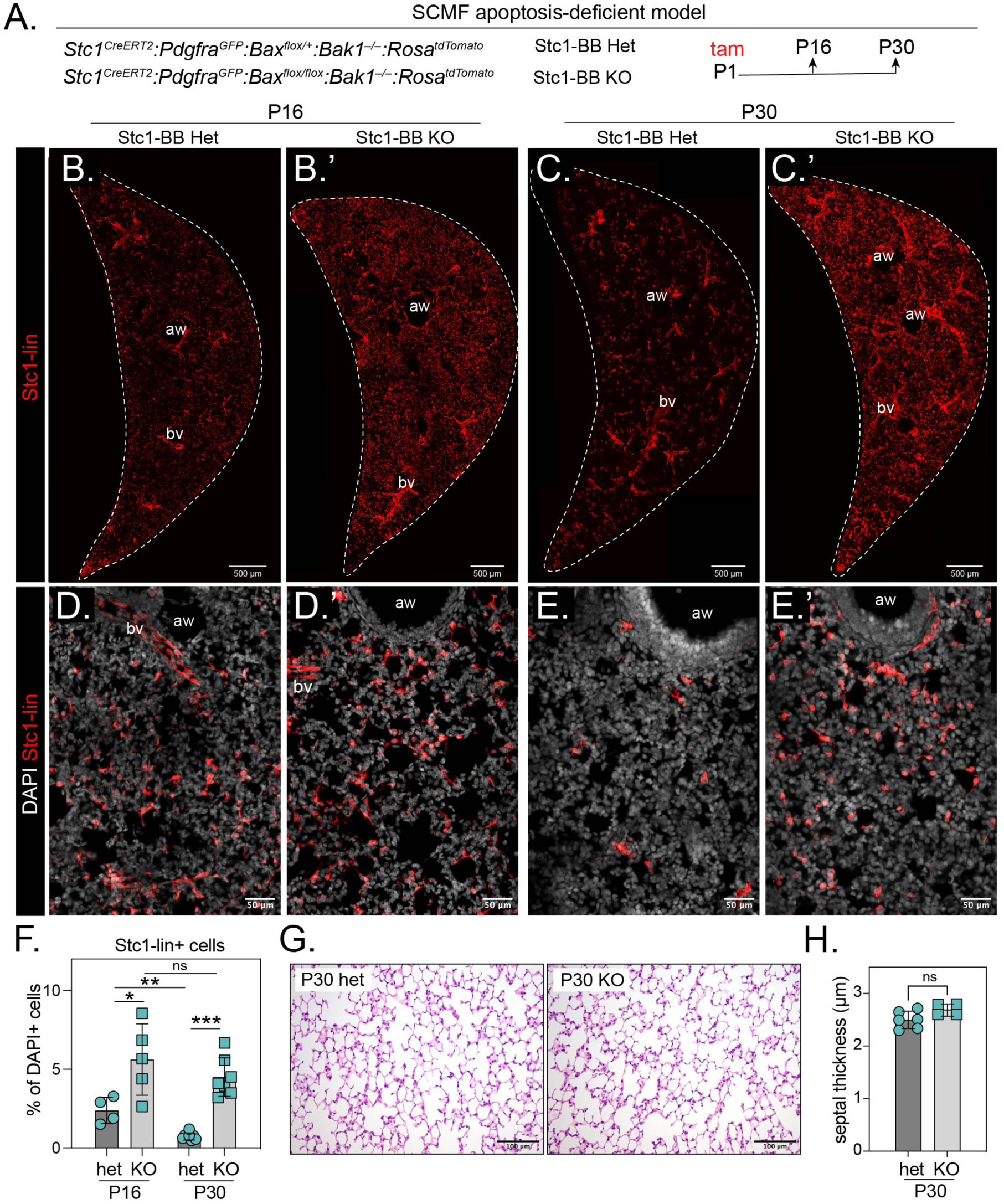
*Stc1*-lineage loss of apoptosis effectors Bax and Bak1 prevents contraction of SCMF population. A) Diagram of *Stc1^CreERT2^*mediated conditional knock out of a floxed *Bax* allele, in a background of *Bak1* knock out, *Pdgfra^GFP^*, and *Rosa^tdTomato^*. B-C)Stc1-BB het (B,C) and Stc1-BB KO (B’, C’) left lung cross sections at P16 (B, B’) and P30 (C, C’). Z-projections of approximately 100 µm. D-E) Higher magnification confocal microscopy of Stc1-BB het (D,E) and Stc1-BB KO (D’, E’) left lung cross sections at P16 (D, D’) and P30 (E, E’). Z-projections of approximately 30 µm. F) Quantification of *Stc1*-lineage in Stc1-BB het and Stc1-BB KO lungs. Blood vessels were excluded from analysis. G) H&E staining of Stc1-BB het and Stc1-BB KO lungs at P30. H) Quantification of alveolar wall thickness from H&E staining in Stc1-BB het and Stc1-BB KO lungs at P30. Abbreviations: Stc1-lin, Stc1-lineage; Tam, tamoxifen; het, heterozygous; KO, knock-out; aw, airway; bv, blood vessel. Scale bars: (B-C’) 500 µm; (D-E’) 50 µm; (G) 100 µm. Graphs represent mean and standard deviation. Statistics: ns, not significant; * *p* < 0.05; ** *p* < 0.01; *** *p* < 0.001.

Because SCMFs are cleared by apoptosis, coinciding with alveolar wall thinning, we hypothesized that preventing SCMF removal would increase alveolar wall thickness. However, by P30, retained Stc1-lineage cells in the Stc1-BB KO appear to have lost their ringlike structure characteristic of SCMFs (Figure 4E’). The alveolar structure appeared identical between control and Stc1-BB KO lungs, and there was no significant difference in alveolar septal thickness (Figure 4G, H). Animal body condition and body weight were unchanged, and we found no significant difference in the abundance of other cell types between Het and KO conditions (Supplementary Figure S4). These data show that SCMFs utilize the apoptotic effectors BAX/BAK1, and that preventing apoptosis does not retain the myofibroblast phenotype of the Stc1-lineage, which become morphologically distinct and have no adverse effect on the alveolar septum.

### “Undead” SCMFs that fail to undergo apoptosis express markers of both SCMF and DMF lineages

To examine the fate of SCMFs prevented from undergoing apoptosis, we performed single-cell RNA sequencing (scRNAseq) on P30 Stc1-BB Het and KO lung mesenchyme (CD45/CD31/Epcam–negative enriched fraction by FACS). We found a cell population that was transcriptomically similar to DMF and ASM (Figure 5A). To test how these Stc1-BB–SCMFs were transcriptomically divergent from normal developmental SCMF, we integrated these data with previously published P7 mesenchyme scRNAseq (Zepp *et al*., 2021). We found that the P30 apoptosis-resistant cells, which we call undead-SCMFs (uSCMF) were distinct from the developmental SCMFs (Supplementary Figure S5A-C). Unlike developmental SCMFs, the uSCMF do not express myofibroblast marker genes, *Acta2* or SM22 protein (Figure 5B, Supplementary Figure S5F). However, the uSCMF expressed *Pdgfra* and *Hhip*, as well as other genes enriched in DMF in our scRNAseq, *Cdh4* and *Igfbp5* (Narvaez Del Pilar *et al*., 2022) (Figure 5B, Supplementary Figure S5D). Other markers were also dysregulated in uSCMF, including some pro-inflammatory genes, *Cxcl14*, *Icam1*, and *Fosb*, although we did not observe any change in immune cell abundance from the H&E stained tissue sections or immunostaining for CD68+ macrophages (Gharib et al., 2009; Lu et al., 2016; Singh et al., 2021) (Supplementary Figures S4C, S4E, S5E and Figure 4G).

**Figure 5.**
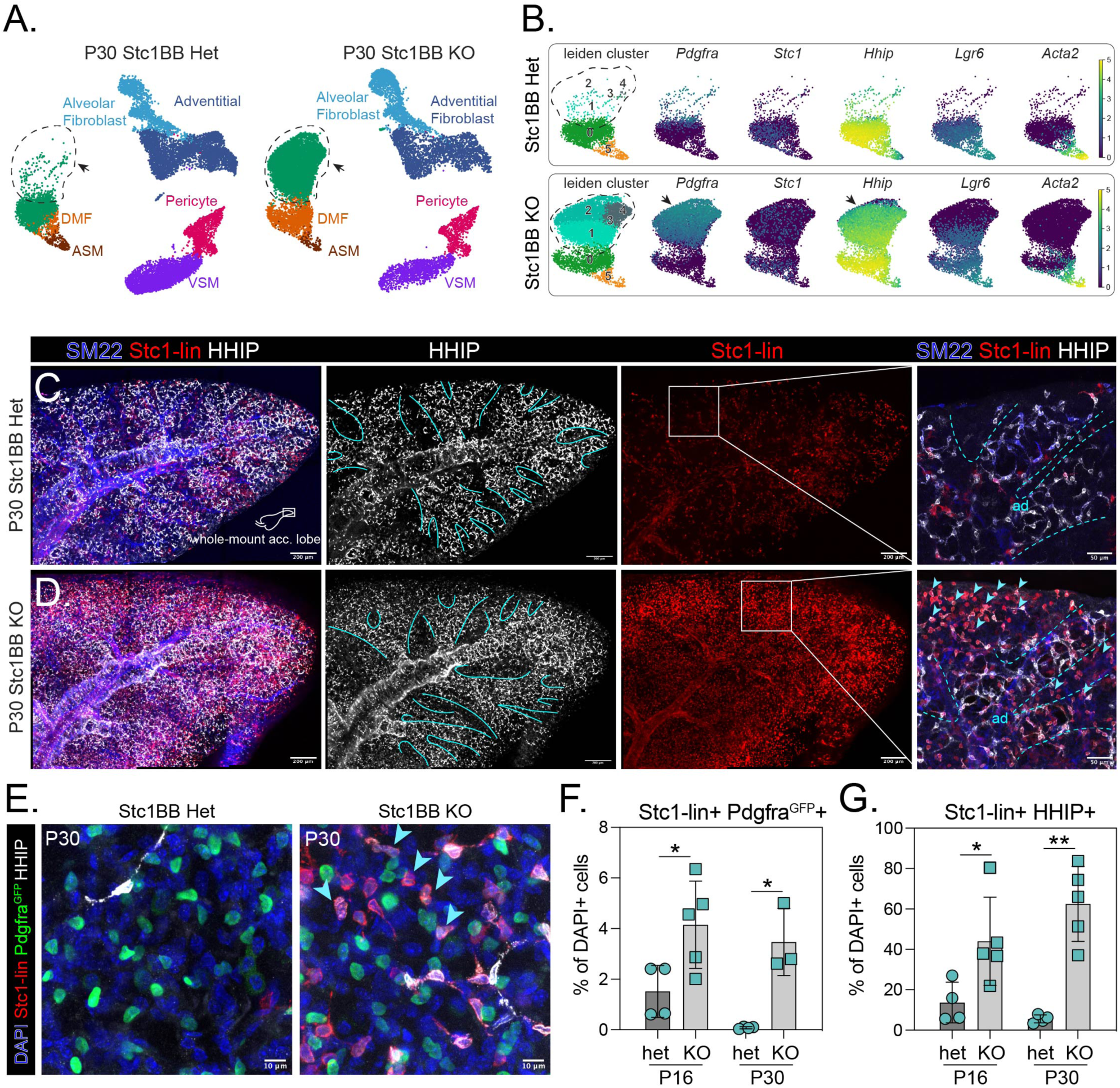
*Stc1*-lineage cells retained in Stc1-BB KO lungs express SCMF and DMF markers but lose myofibroblast identity. A) scRNAseq of P30 mesenchyme in Stc1-BB het and Stc1-BB KO lungs. uSCMF population is outlined. B) Gene expression of SCMF and DMF marker genes in myofibroblast populations in Stc1-BB het and Stc1-BB KO lungs. C-D) Stc1-BB het and Stc1-BB KO whole-mount accessory lobes immunostained for HHIP and SM22. Inset shows HHIP+ SM22+ DMFs, and residual Stc1-lineage cells in the alveolar ducts and alveoli. Z-projections of approximately 250-275 µm (left) and 150 µm (inset). E) Stc1-BB het and Stc1-BB KO lungs expressing Pdgfra^GFP^ and immunostained for HHIP. Arrows indicate triple-positive cells. Z-projections of approximately 80 µm. F-G) Quantification of Stc1-lineage cells expressing Pdgfra^GFP^ (F) and HHIP (G) in Stc1-BB het and Stc1-BB KO lungs at P16 and P30. Abbreviations: Stc1-lin, Stc1-lineage; ASM, airway smooth muscle; DMF, ductal myofibroblast; VSM, vascular smooth muscle; acc. lobe, accessory lobe; ad, alveolar duct. Scale bars: (C, D) 200 µm, inset 50 µm; (E) 10 µm. Graphs represent mean and standard deviation. Statistics: * *p* < 0.05; ** *p* < 0.01.

To investigate the distribution of Stc1-BB KO cells that express HHIP and Pdgfra^GFP^, we performed whole-mount immunostaining and imaging of intact accessory lobes from P30 lungs. Consistent with the scRNAseq results, Stc1-lineage cells in the Stc1-BB KO tissue co-express HHIP and Pdgfra^GFP^ (Figure 5C-G). These data reveal that apoptosis-resistant “undead” SCMFs have dysregulated marker gene expression, including HHIP, and retain Pdgfra^GFP^ expression despite lacking myofibroblast-like morphology.

### Secondary Crest and Alveolar Duct Myofibroblasts develop as distinct lineages

Our results revealed a unique transcriptional state in SCMF or uSCMF, exhibiting a marked increase in HHIP expression at P15 or in P30, respectively (Figures 3, 5). These observations led us to ask whether SCMFs and DMFs are a single developmental lineage or are specified separately. Although early studies speculated a common progenitor for the mesothelium and the SCMF lineage, our prior work with the *Acta2^CreERT2^* localized SCMF progenitors around the distal airspaces (Li *et al*., 2015; Moiseenko *et al*., 2017; Zepp *et al*., 2021). Further, we found that the Stc1-lineage completely excluded WT-1+ mesothelium, thus supporting distinct SCMF and mesothelial lineages (Supplementary Figure S6A,B). To identify the DMF and SCMF progenitors, we performed lineage tracing for these populations starting when SCMFs are specified at embryonic day 15.5 (E15.5) (Zepp *et al*., 2021) (Figure 6A, Supplementary Figure S6C). Like *Hhip, Lgr6* expression marks ASM and DMF in scRNAseq (Narvaez Del Pilar *et al*., 2022) (Figure 3B). Therefore, we also performed Lgr6-lineage tracing using *Lgr6^CreERT2^* to follow the ASM/DMF cells (Lee et al., 2017) (Figure 6A).

**Figure 6.**
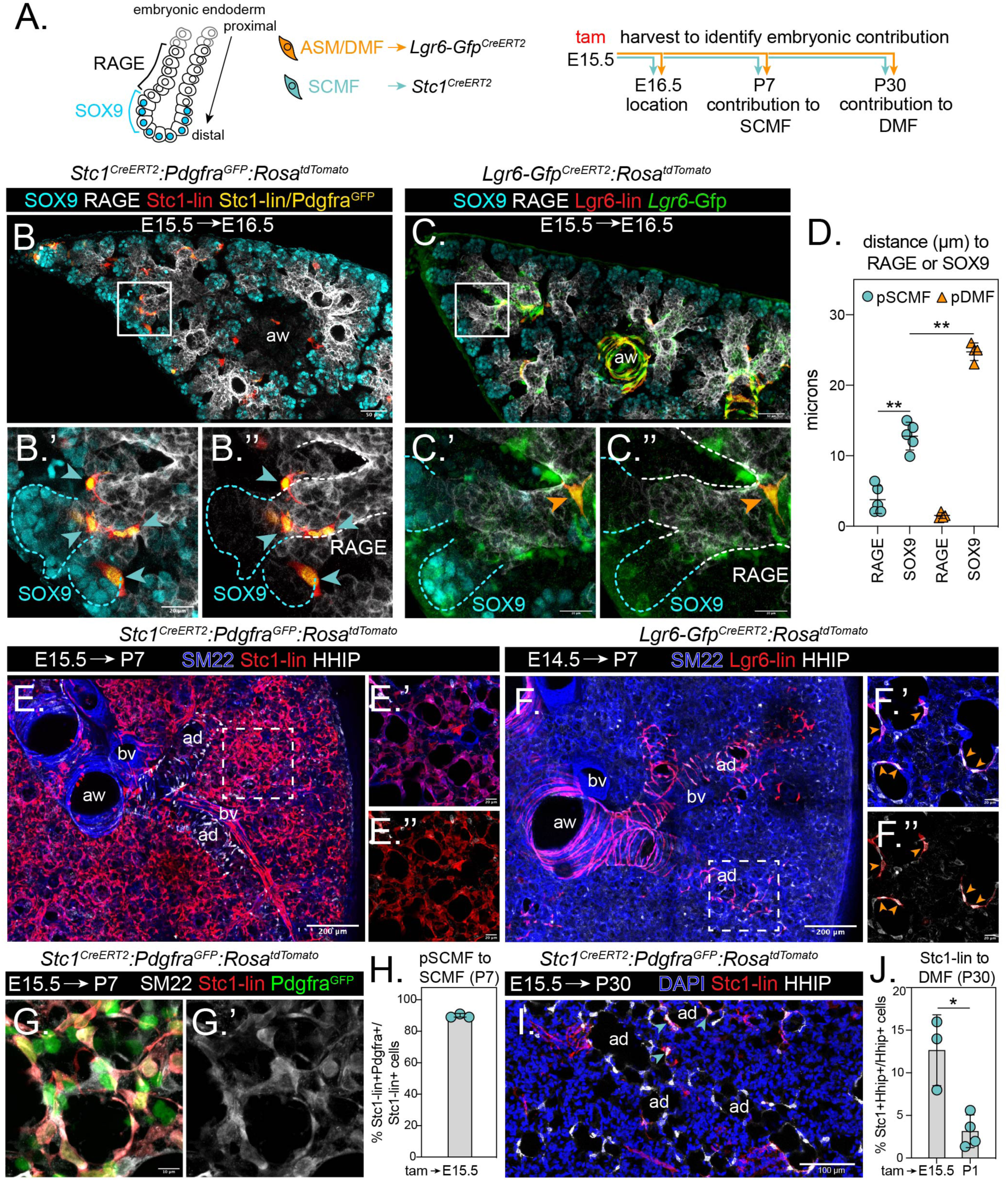
SCMF and DMF are distinct lineages in embryonic development. A) Diagram of proximal to distal embryonic lung endoderm expressing RAGE in the intermediate stalk and SOX9 in the distal tip, and lineage tracing strategy for DMF and SCMF progenitors. B-C) Stc1 (B) or Lgr6 (C) lineage trace from E15.5 to E16.5, immunostained with SOX9 and RAGE. Dashed lines in insets indicate SOX9-expressing distal tip (blue) and RAGE-expressing stalk (white) endoderm, and arrows indicate Stc1-lineage cells expressing Pdgfra^GFP^ (B’, B’’) or Lgr6-lineage cells (C’, C’’). Z-projections of approximately 65 µm (B, B’), 90 µm (C), and 75 µm (C’). D) Quantification of SCMF (B) and DMF (C) progenitors’ distance from the nearest SOX9 or RAGE expressing cell. Data points represent median values of each biological replicate. E-F) Stc1 lineage trace from E15.5 to P7 (E) or Lgr6 lineage trace from E14.5 to P7 (F), immunostained for SM22 and HHIP. Inset (E’, E’’) shows Stc1-lineage contribution to SCMF in alveolar region. Inset (F’, F’’) shows that Lgr6-lineage cells in distal airways co-express HHIP and encircle alveolar ducts (arrows). Z-projections of approximately 35 µm (E, F), 30 µm (E’), and 50 µm (F’). G) Stc1-lineage trace from E15.5 to P7 expressing Pdgfra^GFP^, and co-immunostained with SM22. Z-projections of approximately 30 µm. H) Quantification of Stc1-lineage traced cells from E15.5 to P7 co-expressing Pdgfra^GFP^. I) Stc1 lineage trace from E15.5 to P30 co-immunostained for HHIP. Z-projections of approximately 55 µm. J) Quantification of Stc1-lineage traced cells from E15.5 or P1 to P30 co-expressing Hhip. Note that P1-P30 data is repeated from Figure 3E to illustrate discrepancy with E15.5-P30 data. Abbreviations: Stc1-lin, Stc1-lineage; Lgr6-lin, Lgr6-lineage; tam, tamoxifen; ASM, airway smooth muscle; DMF, ductal myofibroblast; SCMF, secondary crest myofibroblast; pDMF, DMF progenitor; pSCMF, SCMF progenitor; aw, airway; ad, alveolar duct; bv, blood vessel. Data are represented as mean and standard deviation, with individual data points shown. Statistics: * *p* < 0.05; ** *p* < 0.01. Scale bars: (B, C) 50 µm; (B’-C’’) 20 µm; (E, F) 200 µm; (E’-F’’) 20 µm; (G) 10 µm; (I) 100 µm.

First, we lineage labeled SCMF progenitors at E15.5 in the *Stc1^CreERT2:^Pdgfra^GFP^:R26R^tdTomato^*mouse. By E16.5, the SCMF progenitors (Stc1-lineage/Pdgfra^GFP^) were localized to the distal stalk, a transition zone of distal lung endoderm demarcated by the boundary of RAGE-expressing (stalk) and SOX9 (distal tip) epithelial progenitors (Figure 6B, Supplementary Movie 1). In contrast, DMF progenitors labeled by the Lgr6-lineage at E15.5 (excluding airway smooth muscle) were localized to the more proximal RAGE+ stalk epithelium at E16.5 (Figure 6C and Supplementary Movie 2). Measuring the distance between these progenitors and the endoderm revealed that SCMF progenitors are closer to the RAGE+ stalk than SOX9+ distal tips and that DMF progenitors are already significantly farther away than SCMF progenitors from SOX9+ distal tips by the time that SCMFs are specified at E15.5 (Figure 6D).

SCMF and DMF progenitors’ distinct positions suggested lineage divergence by E15.5, and so we subsequently labeled the Lgr6-lineage at E14.5 to capture any potential common progenitor at an earlier stage. While Stc1-lineage+ cells labeled at E15.5 robustly contribute to SCMFs at P7, Lgr6-lineage+ cells labeled as early as E14.5 were spatially restricted to ASM and DMFs at P7 (Figure 6E, F). SCMF identity of Stc1-lineage+ cells labeled at E15.5 was confirmed by colocalization with Pdgfra^GFP^ and SM22 expression (Figure 6G, H). We also looked for contribution to DMFs, which we identified at P30 by HHIP staining. Interestingly, Stc1-lineage contribution to DMF from E15.5 was 13%, significantly more than from P1, which was 3% (Figures 6I, J). These data show that DMF and SCMF progenitors are spatially separated and therefore develop in and are influenced by unique niches.

### Activation of the Hedgehog pathway in Stc1-lineage cells leads to increased SM22 expression in cells that persist post-alveologenesis

The SCMF and DMF are transcriptionally similar, and we have shown that the SCMF expresses DMF marker genes following genetic perturbation (Chaudhry *et al*., 2024) (Figures 1A, 5). While our progenitor lineage tracing demonstrated that DMFs and SCMFs arise from distinct progenitors, we next sought to test whether SCMF fate could be altered by genetically activating pathways that affect myofibroblast development in the lung. We and others previously reported that SCMF progenitors are receptive to Hedgehog (*Shh*) and Wnt signaling from the developing epithelium, and knocking out *Shh* from the distal endoderm at E15.5 inhibited SCMF development (Cohen et al., 2009; Kugler et al., 2017; Zepp *et al*., 2021). We hypothesized that these signaling pathways might inform the segregation of the SCMF and DMF progenitors or affect SCMF apoptosis. To test this hypothesis, we used a Cre-inducible constitutively active *Smoothened homolog* (Hh^GOF^) and stabilized beta-catenin (Wnt^GOF^) to activate the Hh and Wnt pathways, respectively (Harada et al., 1999; Jeong et al., 2004) (Figure 7A, Supplementary Figure S7A).

**Figure 7.**
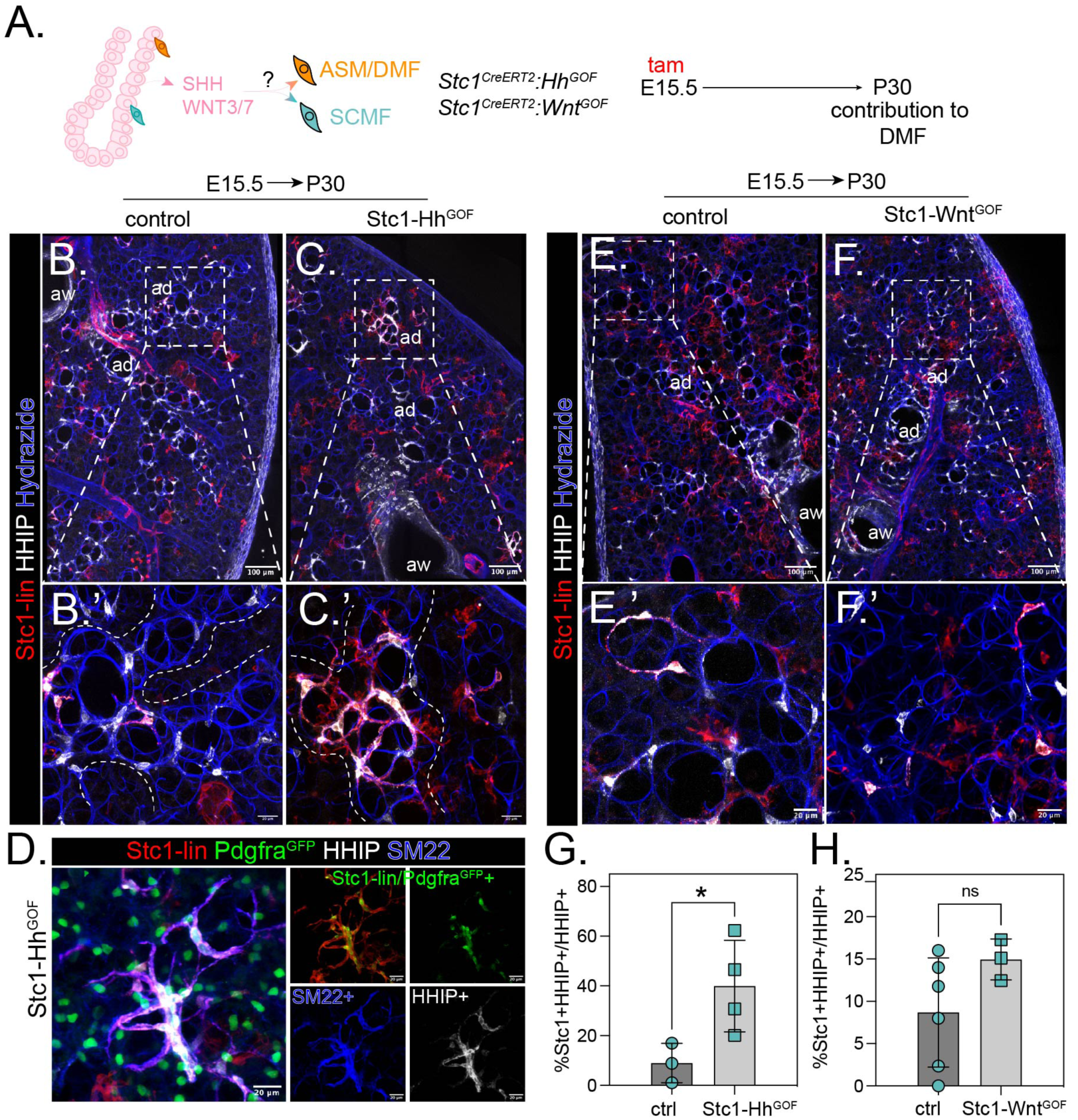
Activation of Hh but not Wnt signaling in SCMF progenitors induces a proximal myofibroblast-like morphology. A) Diagram of genetic strategy to activate Hh and Wnt signaling pathways in SCMF progenitors. B-C) Stc1-lineage traced cells from E15.5 to P30, with (C) or without (B) Hh^GOF^. Insets (B’, C’) of Stc1-lineage traced cells positive for Hhip in distal regions of alveolar ducts. Dashed lines in (B’, C’) outline alveolar ducts. Z-projections of approximately 100 µm (B-C’), 50 µm (E, F), and 130 µm (E’, F’). D) Stc1-lineage traced cells with Hh^GOF^ from E15.5 to P30 and Pdgfra^GFP^, co-immunostained with HHIP and SM22. Stc1-lineage/Pdgfra^GFP^+ is a colocalization channel of *Pdgfra^GFP^*+ nuclei within Stc1-lineage cells. Z-projections of approximately 130 µm. E-F) Stc1-lineage traced cells from E15.5 to P30, with (E) or without (D) Wnt^GOF^. Insets (D’, E’) of Stc1-lineage traced cells positive for Hhip in distal regions of alveolar ducts. G-H) Quantification of Stc1-lineage traced cells positive for Hhip with Hh^GOF^ (G) or Wnt^GOF^ (H). Data are represented as mean and standard deviation, with individual data points shown. Statistics: ns, not significant; * *p* < 0.05. Abbreviations: Stc1-lin, Stc1-lineage; tam, tamoxifen; ASM, airway smooth muscle; DMF, ductal myofibroblast; SCMF, secondary crest myofibroblast; aw, airway; ad, alveolar duct. Scale bars: (B, C, E, F) 100 µm; (B’, C’, D, E’, F’) 20 µm.

We examined the Stc1-lineage at P30 to best detect any lineage conversion or non-apoptotic SCMFs. We found that Stc1-Hh^GOF^ did not prevent the depletion of SCMFs, but Hh activation did induce HHIP expression in the remaining myofibroblasts (Figure 7B, C). These cells appeared to have larger tdTomato+ cell bodies than controls, expressed Pdgfra^GFP^, and also expressed high levels of SM22 (Figure 7D). However, Stc1-Wnt^GOF^ had no significant change to its myofibroblasts or HHIP expressing Stc1-lineaged cells (Figure 7E-H). Further, we activated Hh and Wnt signaling in the Lgr6 lineage at P1 and found no difference between control to SCMF contribution between control, Lgr6–Hh^GOF^, and Lgr6–Wnt^GOF^ lungs at P7 (Supplementary Figure S7A-C). These data suggest that Hh pathway gain of function in SCMF results in dysregulated myofibroblast identity but is insufficient to expand SCMFs or alter their lineage potential.

## DISCUSSION

In this study, we generated a knock-in *Stc1^CreERT2^* mouse and validated that it efficiently labels SCMFs. We analyzed the clonal expansion and contraction by apoptosis of Stc1-lineage SCMFs. Notably, Stc1-lineage does not label DMFs, although we found that the DMF marker Hhip has transient overlap with SCMFs near the end of their life. Genetically deleting SCMFs’ ability to undergo apoptosis resulted in residual ‘undead’ Stc1-lineage cells that lost their myofibroblast identity but expressed Pdgfra^GFP^ and HHIP. We investigated the embryonic origins of SCMF and DMF lineages by lineage tracing, finding them separate by the time *Stc1*+ SCMFs are specified at E15.5. In genetic gain-of-function experiments of either Hh and Wnt signaling pathways, which are known for controlling myofibroblast development, cell fate or positional identity in SCMF were not broadly changed. Overall, the *Stc1^CreERT2^* model illuminates neonatal myofibroblast lineages and their developmental origins, which can help resolve the cellular and molecular dynamics of lung development and disease.

The *Stc1^CreERT2^* improves on prior lineage tracing tools used to describe SCMFs, which were limited in their specificity to distinguish SCMFs from other mesenchymal cell types. DMFs were recently described as a separate population of myofibroblasts (Chaudhry *et al*., 2024; Khan et al., 2024; Narvaez Del Pilar *et al*., 2022; Yie et al., 2023), and conclusions about *Acta2*-expressing neonatal alveolar fibroblasts could pertain to either DMFs or SCMFs. Recent studies have suggested that ‘dedifferentiated’ Acta2-lineage cells that persist post-alveologenesis and lose Acta2 expression re-differentiate into myofibroblasts in adult injury models, including hypoxia, bleomycin, and influenza (Chandran et al., 2024; Khadim et al., 2025). *Gli1^CreERT2^* labels multiple mesenchymal lineages, including mesothelium and SMA-negative lipofibroblasts, and myofibroblasts analyzed at P14 are likely enriched for DMFs as SCMFs undergo apoptosis (Hagan *et al*., 2020; Li *et al*., 2015). *Fgf18^CreERT2^* labels a subset of *Gli1*+ mesenchyme that is developmentally cleared, but also labels mesothelium, peribronchiolar smooth muscle, and 23% of AT1s with postnatal labeling (Hagan *et al*., 2020). The *Pdgfra^rtTA^*-lineage contributes to both SCMFs and DMFs, with scRNAseq data capturing clusters expressing DMF markers *Hhip* and *Lgr6* (Li *et al*., 2018). Therefore, future studies on neonatal-derived myofibroblasts must incorporate specific markers and imaging that can distinguish between ASM, DMF, and SCMF.

Stc1-lineage labeling allows us to distinguish SCMFs from DMFs and study them separately. We have demonstrated that SCMFs and DMFs have unique positional and molecular profiles in the postnatal lung and are specified in spatially distinct compartments. The *Stc1*+ SCMF progenitors are at the RAGE-SOX9 transition zone, whereas Lgr6-lineage DMF progenitors arise more proximally along the RAGE+ epithelium, farther away from the SOX9+ distal tips. Both are likely to be in the HOPX+ stalk and a recently described ICAM+ transition zone (Frank et al., 2019; Ke et al., 2025). ASM, DMFs, and SCMFs exist in a proximal-distal axis reflected by a gradient of Hh signaling activity (Liu et al., 2013). The more proximal smooth muscle and DMF populations express HHIP, which binds to the Hh ligand and inhibits Hh signaling (Chuang et al., 2003; Chuang and McMahon, 1999), while the distal SCMFs are HHIP-negative and GLI1-positive, indicators of active Hh signaling (Kugler *et al*., 2017; Li *et al*., 2015; Liu *et al*., 2013). SCMFs transiently express HHIP prior to apoptosis, and residual SCMFs that could not undergo apoptosis continue to express HHIP, suggesting that HHIP and/or Hh signaling may maintain SCMF survival or affect SCMF apoptosis (Kugler *et al*., 2017).

We found that SCMFs expand clonally from *Stc1*+ progenitors. SCMFs are highly proliferative relative to other populations as SCMFs expand postnatally, and SCMF clones populate neighboring alveoli. Stc1-lineage embryonic SCMF progenitors specified between E14.5 and E15.5 give rise to most SCMFs. However, more progenitors are specified by P1, presumably due to further branching of the lung between E15.5 and P1 that gives rise to new SCMF progenitor niches at newly formed stalk-tip boundaries (Alanis et al., 2014; Frank *et al*., 2019; Ke *et al*., 2025). The decrease in Stc1-lineage cell number between P10 and P15 corresponds with the timing of increased Stc1-lineage positivity for cleaved CASPASE 3. This is in agreement with our prior observations that SCMFs have higher mRNA expression of apoptosis pathway components at P7 (Chaudhry *et al*., 2024; Zepp *et al*., 2021), and other studies examining apoptosis kinetics in Pdgfra-lineage or Acta2-lineage populations (Chandran *et al*., 2024; Narvaez Del Pilar *et al*., 2022). CASPASE 3 cleavage can indicate either intrinsic apoptosis (caused by factors including growth factor depletion, ER stress, replication stress, or changes in cellular attachment signaling such as integrin or Hippo signaling) or extrinsic apoptosis (caused by death receptor activation) (reviewed in Yuan *et al*. (Yuan and Ofengeim, 2024)). Stc1-lineage SCMFs deficient in intrinsic apoptosis effectors BAX and BAK1 persist past the apoptotic period, suggesting that SCMFs are eliminated by intrinsic apoptosis. Together, these data further detail the transient life cycle of SCMFs, adding cell-type specificity to prior studies of expansion and apoptosis, comparison of cellular proliferation rates during bulk alveologenesis, clonal expansion from progenitors, and support for an intrinsic model of SCMF apoptosis (Bruce *et al*., 1999; Gao et al., 2022; He et al., 2021; Khan *et al*., 2024; Kimani et al., 2009; Li *et al*., 2018; McGowan and McCoy, 2015; Negretti *et al*., 2021; Schittny *et al*., 1998).

We hypothesized that Hh signaling, which is necessary in the Acta2-lineage for alveolar development (Kugler *et al*., 2017; Zepp *et al*., 2021), imposes the proximal-distal positional identity of DMF and SCMF and that Hh signaling loss induces apoptosis. However, enforced Hh signaling in the Stc1-lineage imposed a muscularized phenotype, with retained Stc1-lineage cells exhibiting a smooth muscle-like morphology. Wnt signaling, which is present in alveolar myofibroblast development and is critical for smooth muscle development (Cohen *et al*., 2009; Fang et al., 2022; Zepp *et al*., 2021), did not affect SCMF or DMF lineages when constitutively active. These data show that SCMFs and DMFs are molecularly and spatially distinct lineages from the time of *Stc1*-expressing SCMF-progenitors, which are mostly lineage committed and unchanged by enhancing Hh or Wnt signaling pathways. Future studies targeting additional pathways and extensive embryonic induction timepoints could identify critical factors and specific times that regulate lineage plasticity. A revised catalog of heterogenous fibroblast lineages during lung development and in disease will require specific lineage tracing tools to demonstrate plasticity between fibroblast types and differentiation. Due to its clonal proliferation, alveolar localization, and critical function, future studies can employ the *Stc1^CreERT2^*mouse model to address the involvement of SCMF in pediatric lung disease or outcomes of early life exposures on SCMF activity and alveolar maturation.

### Limitations

A limitation of using *Stc1^CreERT2^* to study SCMFs is that *Stc1* is also expressed in a subset of pericytes and arterial endothelium. Pdgfra^GFP^ was used in our study as an additional marker of SCMFs when morphology alone could not be used to exclude these populations. An intersectional genetic approach would be needed to ablate the Stc1-lineage without damaging arterial endothelium. A second limitation is that while HHIP is a DMF marker post-alveologenesis, we find that some SCMFs transiently express HHIP as they near apoptosis, and S*tc1-*BB KO ‘undead’ SCMFs also express HHIP. In this case, HHIP may represent a cell state of reciprocal Hh-HHIP regulation, rather than a cell fate. However, our use of *Lgr6^CreERT2^*provides an independent line of evidence for our conclusions about DMFs.

## METHODS

### Mice

#### Generation of the Stc1-CreERT2 knock-in allele

A targeting construct sequence generously provided by Dr. Mark Kahn included a T2A self-cleaving peptide sequence, CreERT2 coding region, and a FRT-flanked neomycin resistance cassette. 1 kb arms of homology were added to target this construct to replace the *Stc1* stop codon. Two guide RNA sequences targeting the 3’ UTR near the stop codon (5’-ATGCAATTTTTCTTTAACGG-3’ and 5’-CCCCTAAAATGCTATTAGTT-3’) were cloned individually into pX330 (Addgene, plasmid #42230) and the corresponding PAM NGG sites were mutated on the targeting construct. The targeting construct was synthesized by Vectorbuilder.

Genetic targeting of V6.5 mouse embryonic stem (ES) cells used to generate the Stc1-CreERT2 knock-in allele was previously described (Chen et al., 2022). Following G418 selection, individual clones were screened by PCR for correct insertion at the 5’ (primers 5’-ACCCCAAGATGTAGGGGAGC −3’ and 5’-CACGTCCCCGCATGTTAGAAGA −3’) and 3’ ends (primers 5’-CTTGGCGCTACACAAGTGGC −3’ and 5’-AGCCCAAGTTTCCCGGACTA −3’). PCR amplicons were verified by Sanger sequencing.

Generation of chimeras by ES cell microinjection into blastocysts was performed by the Transgenic Mouse Core at the University of Pennsylvania School of Veterinary Medicine as previously described (Hong et al., 2020). Chimeras were screened by PCR with the 5’ and 3’ targeting primers, and with genotyping primers (5’-AGCCTGATGGAGAAGATCGGG −3’ and 5’-TGACTTCATCAGAGGTGGCATCC −3’). Chimeras were bred to *Actb-Flpe* mice (The Jackson Laboratory, stock no. 005703) to remove the FRT-flanked neomycin resistance cassette, and offspring were screened by PCR for its removal (5’-CGCTACTTCTAGTAGGTTCCGC −3’ and 5’-CTGTGAATACCTCTCCCTGC −3’) and then outbred to remove the *Actb-Flpe* gene. *Stc1-CreERT2* positive and *Flpe* negative mice were used in this study.

#### Mouse strains

The following mice were obtained from the Jackson Laboratory: Lgr6EGFP-Ires-CreERT2 (stock no. 016934) (Snippert et al., 2010a), PdgfraH2B:eGFP (stock no. 007669) (Hamilton *et al*., 2003), R26RtdTomato (stock no. 007914) (Madisen *et al*., 2010), R26RConfetti (stock no. 017492) (Snippert *et al*., 2010b), R26RSmoM2 (stock no. 005130) (Jeong *et al*., 2004), and Baxflox;Bak1KO (stock no. 006329) (Takeuchi *et al*., 2005). Ctnnb1 exon 3 floxed mice are previously described (Harada *et al*., 1999). Deletion of floxed Bax alleles was confirmed by two primer pairs spanning the deleted region: 5’-CGCACGTCCACGATCAGTC −3’ and 5’-GAGGAAGTCCAGTGTCCAGC −3’, and 5’-CCTGCAGCGAGCGATGAT −3’ and 5’-CACGGAGGAAGTCCAGTGTC −3’.

#### Tamoxifen and EdU dosing

Tamoxifen (Toronto Research Chemicals, cat. # T006000) was dissolved at 5mg/mL in corn oil (cat. # C8267, Millipore Sigma) and 10% ethanol (Pharmco, cat. # 111000200). Neonatal administration was performed on postnatal day 1 (P1) at a dose of 50 mg/kg by intraperitoneal injection. Adult administration was performed at 8 weeks at 200 mg/kg by oral gavage. Embryonic administration was performed at E15.5 or E17.5 by oral gavage of the dam at a dose of 200 mg/kg tamoxifen and 100 mg/kg progesterone from a stock of 20 mg/mL tamoxifen and 10mg/mL progesterone (Millipore Sigma, cat. # M1629).

EdU (Thermo Fisher, cat. # E10187) was dissolved at 5 mg/mL in PBS and injected intraperitoneally at a dose of 50 mg/kg, 4 hours prior to harvest.

### Histology

Lungs were harvested and thick-cut lung slices were prepared using a vibratome for whole-mount immunofluorescence as previously described (Zepp et al., 2017), and described in brief below.

Mice were euthanized by CO2 inhalation and cervical dislocation, and pups aged P10 or younger were also anesthetized on ice. Lung vasculature was cleared of blood by injecting PBS into the right ventricle. Lungs were inflation-fixed by instilling 2% paraformaldehyde (PFA) at constant pressure from a height of 15cm (P3-P16) or 35cm (P30 and adult), and fixed overnight in PFA. E16.5 and E18.5 lungs were fixed by overnight immersion in 2% PFA.

Left lungs were vibratome sectioned at 150um, permeabilized with 1% Triton X-100 in PBS, and immunostained. All tdTomato and GFP fluorescence was amplified by immunostaining. Fluorescently labeled hydrazides were used to label elastin. All antibodies and hydrazides are listed in Supplementary Table 1. EdU incorporation was visualized using a two-hour reaction with Click-It Plus EdU Alexa Fluor 647 (Thermo Fisher, cat. # C10634) prior to antibody staining. 150µm sections were optically cleared using Scale A2 (Hama et al., 2011). Whole accessory lobes were immunostained and cleared using the CUBIC protocol (Susaki et al., 2015). H&E staining was performed on 6 µm sections from fixed tissue dehydrated by an ethanol gradient and embedded in paraffin.

### Imaging

Imaging was performed on a Leica SP8 confocal microscope (immunofluorescence) or a Leica DMi8 Thunder microscope (H&E) system. Images were processed and analyzed using Imaris (Oxford Instruments) and FIJI (Schindelin et al., 2012). Some images had the following functions performed in FIJI: despeckle, remove outliers, and clear outside (empty space surrounding lung). Co-expression channels of Stc1-lineage and Pdgfra^GFP^ were created with FIJI’s Image Calculator “AND” function on each z-plane prior to z-projection. Supplemental movies were created in Imaris and edited in Adobe Premier Pro. Cell counts were measured with the “spots” function in Imaris from Z-projections acquired with a 40x objective at 1 µm z-step intervals. Cellular distances in Figure 6 were measured using the “distance transformations” function in Imaris.

Alveolar septal thickness was measured by manually drawing lines in FIJI perpendicularly across alveolar walls, using high-resolution images of H&E staining in regions lacking large airways and blood vessels. Three images from multiple sections per animal were used to generate a minimum of 60 measurements per animal.

### Flow cytometry

Lungs were prepared as a single-cell suspension for flow cytometry as previously described (Zepp *et al*., 2017). Briefly, lungs were minced and digested in a mixture of 480U/mL Collagenase Type I (Gibco, cat. # 17100017), 50U/mL Dispase (Corning, cat. # 354235), and 0.33U/mL DNase I (Roche, cat. # 10104159001) at 37C with agitation for 20-45 minutes. The cell pellet was treated with ACK buffer to lyse red blood cells, strained through 100µm and 40µm cell filters, and incubated for 10 minutes at room temperature with flow cytometry antibodies listed in Supplementary Table 1. Cells were resuspended in FACS buffer (2% FBS, 2 mM EDTA, 25 µm HEPES, and 1x Penicillin/Streptomycin in HBSS) containing DAPI (2.5 ug/mL) as a live/dead stain, and analyzed on a LSR Fortessa (BD Biosciences) and FlowJo software.

### Statistical analysis

GraphPad Prism was used for graphing and statistical analysis. Mann-Whitney tests were used for all pairwise comparisons. P-values less than 0.05 were considered significant. Multiple comparisons were analyzed by Kruskal-Wallis tests, and the significance threshold of post-hoc comparisons was adjusted for the number of comparisons (*p* < 0.05/n).

### Single Cell RNA Sequencing

Single cell suspensions from lungs were prepared as above. Cells from individual animals (3 per genotype: 2 male and 1 female) were labeled with unique Cell Multiplexing Oligos (10x Genomics 3’ CellPlex Kit Set A, cat. # 1000261) according to the Chromium Single Cell 3’ v3.1 Cell Multiplexing Protocol 4, modified by using non-barcoded, fluorescently labeled antibodies for enrichment of mesenchyme. Individuals were pooled into two samples by genotype (Stc1-BB het and KO), and negatively sorted for EPCAM (epithelium), CD31 (endothelium), and CD45 (immune) by a BD FACSAria II Cell Sorter. An aliquot of cells from each animal was reserved to confirm by FACS the expected phenotype of ectopic tdTomato+ mesenchyme in each Stc1-BB KO and none of the Stc1-BB het animals.

Sorted cells were counted by Trypan Blue exclusion and resuspended according to the 10x Genomics Protocol for a target recovery of 30,000 cells per genotype. Single cell suspensions were loaded into a 10x Genomics Chromium Controller, and separate libraries from each genotype were generated according to the Chromium Single Cell 3’ v3.1 Cell Multiplexing Dual Index Library protocol.

### Bioinformatics

scRNA-seq data generated in this study were processed as follows. Reads in FASTQ format were aligned to the NCBI RefSeq GRCm39 genome, and reads with unique molecular identifiers (UMIs) were quantified at the gene level using STAR-solo, with *soloUMIdedup=”1MM_CR”*, *soloUMIfiltering=”MultiGeneUMI_CR”,* and *soloCellFilter=”EmptyDrops_CR”* (Dobin et al., 2013). Ambient RNA was removed from the generated counts matrices using the scvi-tools implementation of scAR (Sheng et al., 2022; Xu et al., 2021). Ambient-corrected counts matrices were subsequently processed using scanpy and anndata (Wolf et al., 2018). Cells were filtered to include only those cells with more than 500 unique genes detected/cell, fewer than 6581 unique genes detected/cell (the 95th percentile), and fewer than 15% of UMIs attributed to mitochondrial genes. Genes were filtered from the combined dataset if detected in fewer than 5 cells. Putative cell doublets were identified and removed from each batch independently using the scvi-tools implementation of the SOLO doublet identification model (Bernstein et al., 2020). Cell hashing oligo demultiplexing was performed using 10X Genomics Cell Ranger v7.1.0 to assign biological replicate IDs to cells for each genotype.

Gene expression was calculated from scAR-corrected UMI-counts by first normalizing to counts per 10k UMIs (CP10k) using the *sc.pp.normalize_total* function with *target_sum=10000* and then calculating ln(CP10k + 1) using the *sc.pp.log1p* function. The top 2000 highly variable genes in the dataset were identified using the *sc.pp.highly_variable_genes* function, with *flavor=”seurat_v3”* and *n_top_genes=2000*. An initial principal component analysis was performed using the *sc.tl.pca* function with *svd_solver=”arpack”*. This PCA was subsequently batch-corrected using the pytorch implementation of the harmony batch correction algorithm (Korsunsky et al., 2019). A cell-cell neighborhood was calculated using the *sc.pp.neighbors* function, with *n_neighbors=int(0.25 * sqrt(adata.n_obs))*, and this cell-cell neighborhood was used to generate a 2-dimensional UMAP using the *sc.tl.umap* function with *init_pos=”spectral”* (McInnes et al., 2020). Finally, cell clustering was performed using the leiden algorithm via the *sc.tl.leiden* function (Traag et al., 2019).

Testing for differential gene expression was performed using the *scanpy st.tl.rank_genes_groups* function with *test=’wilcoxon’*. Over-representation analysis (ORA) was performed on genes with resulting FDR < 0.05 using gProfiler, with runtime options *ordered query*, *only annotated genes*, *g:SCS threshold<0.05*, and *max term size=5000* (Kolberg et al., 2023).

## ACKNOWLEDGEMENTS

We thank Mark Kahn’s laboratory at the University of Pennsylvania for assistance with designing the *Stc1^CreERT2^* construct and for advice on gene targeting. We gratefully acknowledge the contributions of N. Adrian Leu and the University of Pennsylvania School of Veterinary Medicine Transgenic Mouse Core for mouse ES cell blastocyst microinjection services, the CHOP Flow Cytometry Core Laboratory, and the CHOP Center for Applied Genomics Sequencing Core.

## FUNDING

This work was supported by the National Institutes of Health (R00HL141684, R35GM119461) to J.A.Z., (K08HL150226) to J.B.K, (R56HL167937) to D.B.F. We are grateful for the generous support provided by the Ayla Gunner Prushansky Research Fund.

## CONFLICTS OF INTEREST

The authors declare no competing or financial interests.

---

**Supplementary Figure 1.**
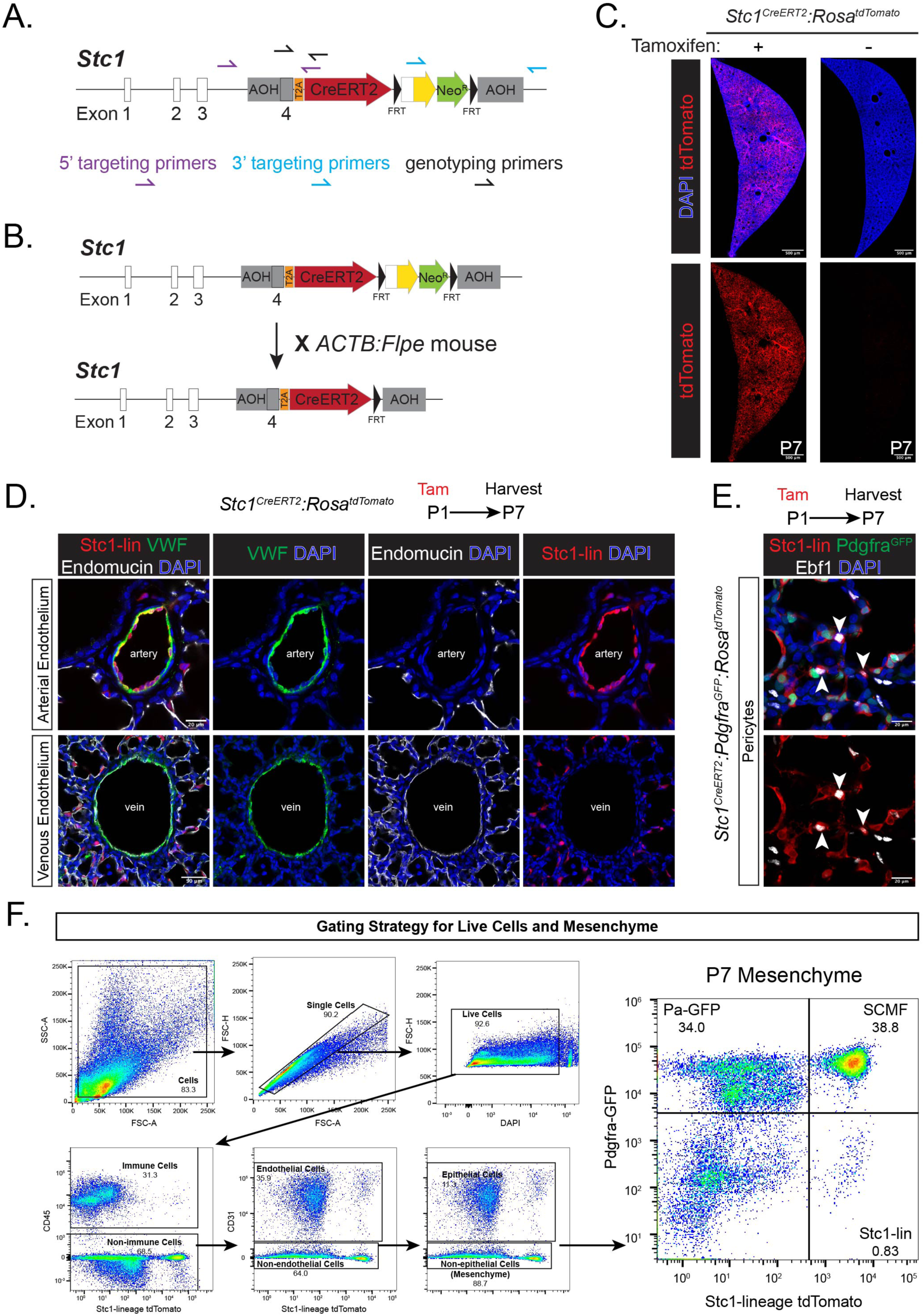
*Stc1^CreERT2^* non-SCMF targets include arterial endothelium and pericytes. A) Design of *Stc1^CreERT2^*mouse. 5’ and 3’ targeting primer pairs binding outside of the arms of homology were used to confirm correct insertion by PCR and Sanger sequencing. Genotyping primers within the arm of homology and CreERT2 sequence were used to confirm genotype in subsequent generations. B) Strategy to remove the FRT-flanked *Neo^R^*cassette by crossing to the *ACTB*:*Flpe* mouse. C) *Stc1^CreERT2^*:*R26R^tdTomato^* left lung cross-section at P7 with or without tamoxifen administration at P1. Z-projections of approximately 100 µm. D) *Stc1^CreERT2^*:*R26R^tdTomato^* lungs at P7 immunostained for VWF and Endomucin. Lumens of VWF+ macrovessels are labeled artery (top row) or vein (bottom) based on the respective absence or presence of Endomucin. Single Z-planes shown. E) *Stc1^CreERT2^*:*Pdgfra^GFP^:R26R^tdTomato^*lungs at P7, immunostained for Ebf1. Arrowheads indicate Stc1-lineage+Ebf1+ pericytes. Z-projections of approximately 5 µm. F) Gating strategy for the flow cytometry of all live single cells (Figure 2F,G) and all mesenchyme (Figure 2H,I). Abbreviations: Stc1-lin, Stc1-lineage; Pa-GFP, Pdgfra^GFP^; tam, tamoxifen. Scale bars: (C) 500 µm, (D) top row 20 µm, bottom row 50 µm, (E) 20 µm.

**Supplementary Figure 2.**
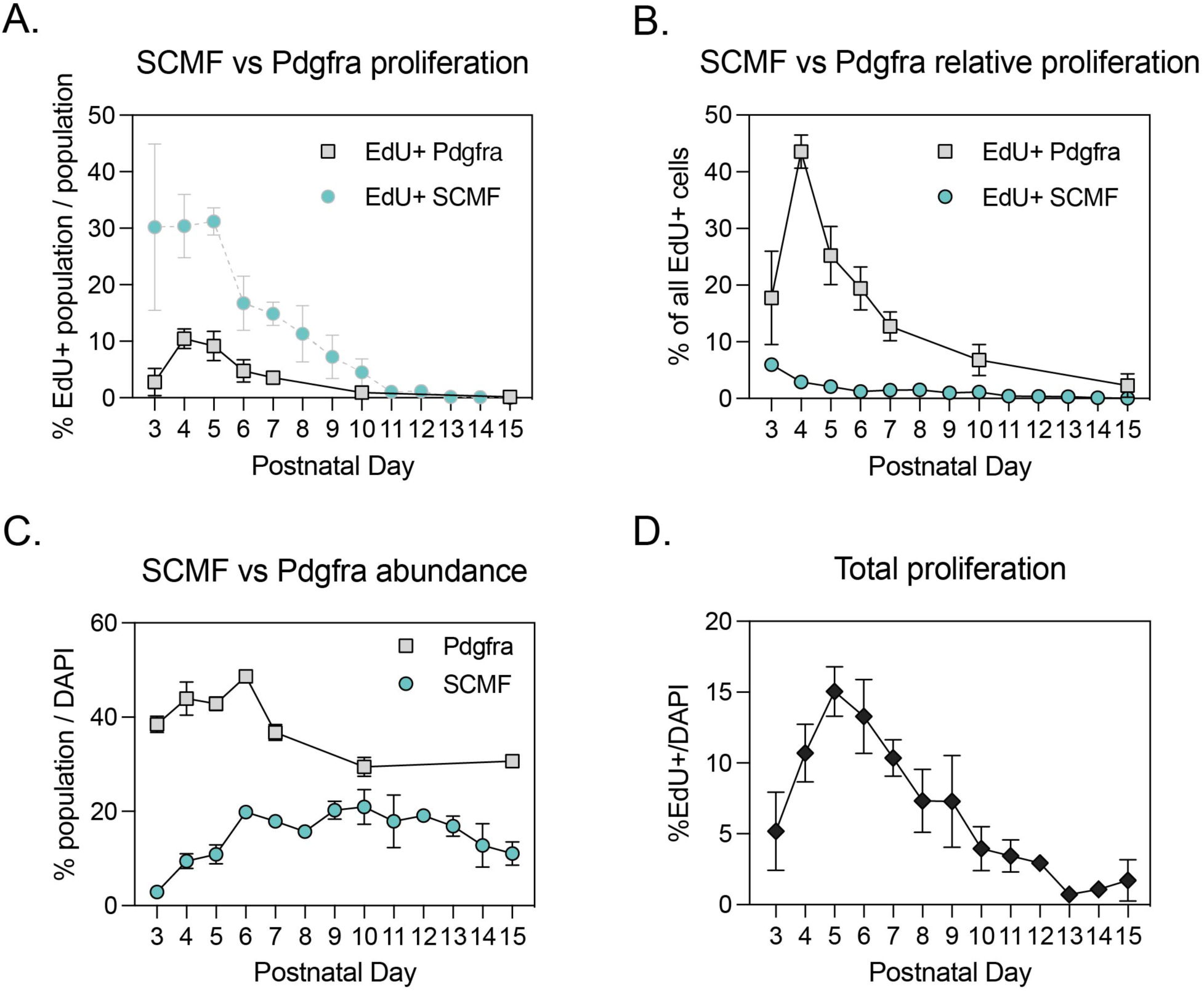
Proliferation of SCMF peaks earlier than non-SCMF Pdgfra+ fibroblasts and total lung cells. A) Percentage of SCMF (Stc1-lineage+) or Pdgfra (Stc1-lineage negative) cells that are EdU+ at the indicated postnatal timepoints during alveologenesis. Note that these SCMF data are repeated from Figure 2F, for illustrative purposes to compare to Pdgfra cells. Tamoxifen was administered at P1 for all data in Supplementary Figures S2A-D. B) Percentage of all EdU+ cells that are SCMF or Pdgfra. C) Percentage of all DAPI+ cells that are SCMF or Pdgfra. D) Percentage of all DAPI+ cells that are EdU+. Data are shown as mean and standard deviation.

**Supplementary Figure 3.**
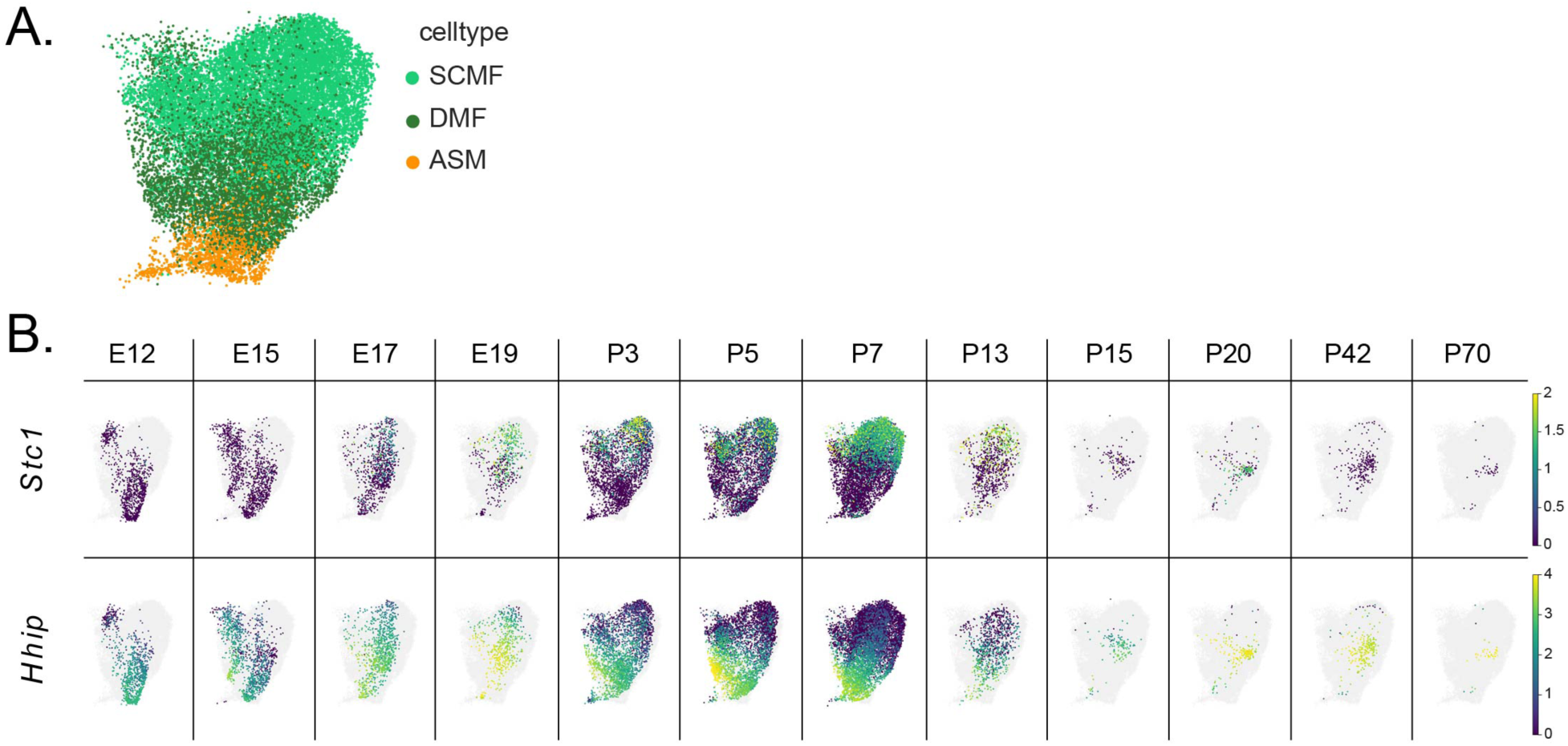
*Stc1* and *Hhip*-expressing populations are mostly distinct during lung development. A) UMAP of scRNA-seq showing myofibroblast populations throughout lung development, subset from Figure 1A. B) Comparison of *Stc1* and *Hhip* expression throughout lung development.

**Supplementary Figure 4.**
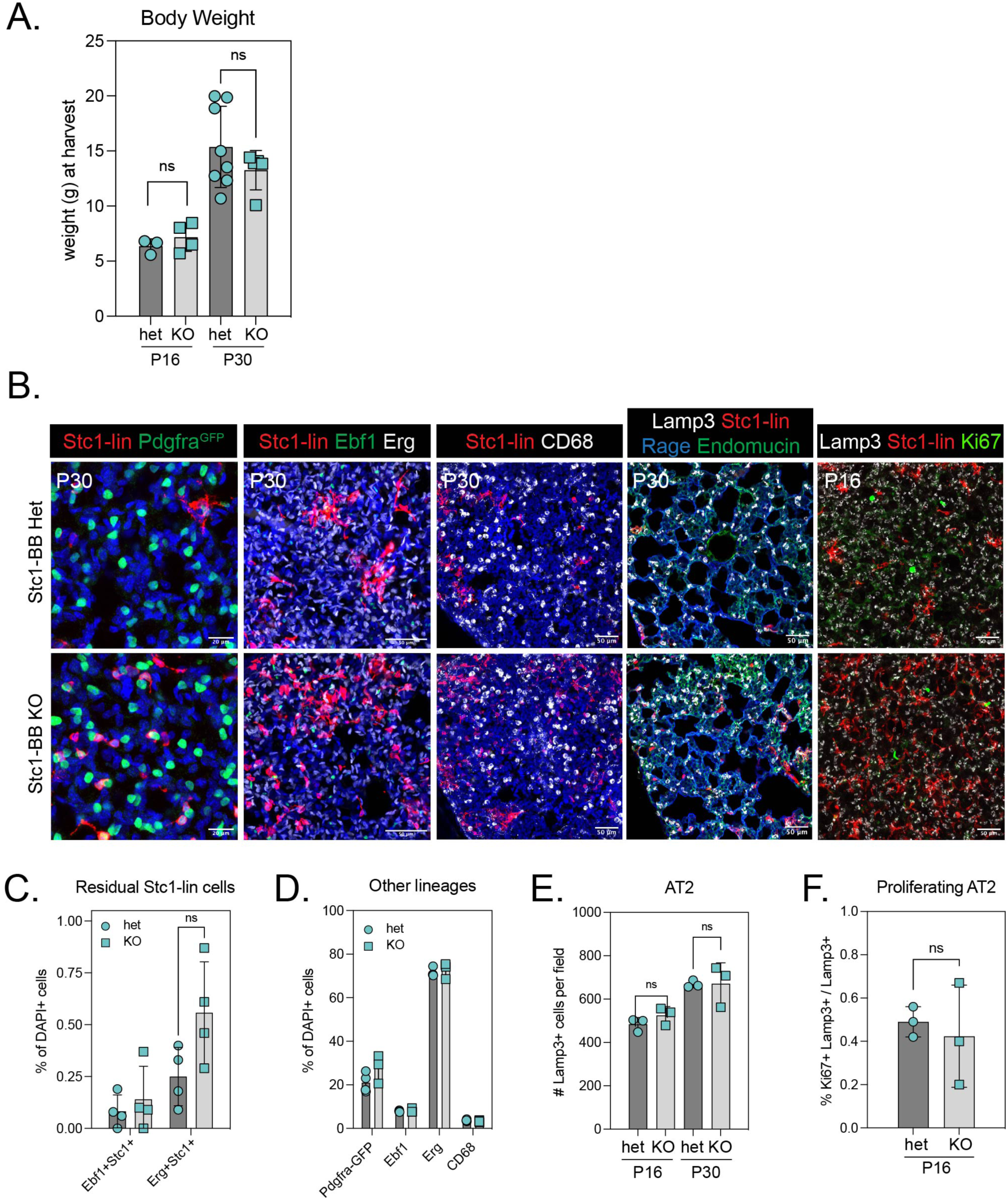
*Stc1*-lineage loss of apoptosis effectors Bax and Bak1 does not impact abundance of other lung lineages. A) Body weights of Stc1-BB het and Stc1-BB KO mice at P16 and P30. B) Stc1-BB het and Stc1-BB KO lungs immunostained for markers of various lung cell types at P30, except the right column at P16. Z-projections of approximately (left to right) 50 µm, 80 µm, 100 µm, 20 µm, 100 µm. C) Quantification of Stc1-lineage cells in Stc1-BB het and Stc1-BB KO lungs at P30 co-expressing pericyte marker Ebf1 or endothelium marker Erg. D) Quantification of cell lineages in Stc1-BB het and Stc1-BB KO lungs at P30, irrespective of Stc1-lineage positivity, including Pdgfra^GFP^+ alveolar fibroblasts, Ebf1+ pericytes, Erg+ endothelium, and CD68+ macrophages. E-F) Quantification of total Lamp3+ AT2 cells (F) or Ki67+Lamp3+ proliferating AT2 cells (F) per field of alveolar space in Stc1-BB het and Stc1-BB KO lungs. Fields were of equal size for all data points: 388.26 µm (x) x 388.26 µm (y) × 100 µm (z). Scale bars: (B) left column, 20 µm; other columns, 50 µm. Abbreviations: Stc1-lin, Stc1-lineage; WT, wild-type; het, heterozygous; KO, knock out; gDNA, genomic DNA; ns, not significant; AT2, alveolar epithelial cell type 2. Data are represented as mean and standard deviation, with individual data points shown. Statistics: ns, not significant.

**Supplementary Figure 5.**
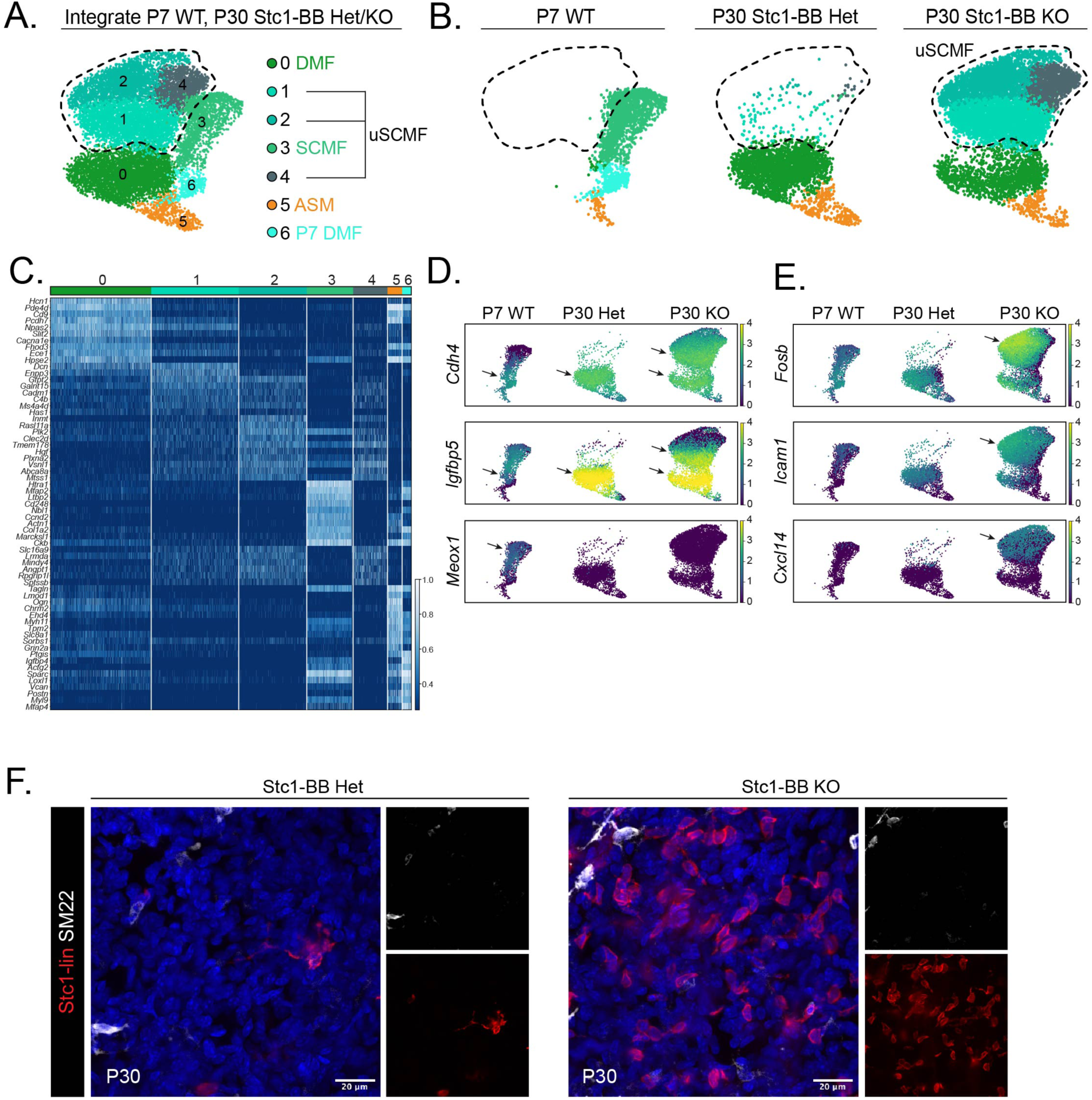
*Stc1*-lineage cells retained in Stc1-BB KO lungs have altered gene expression compared to P7 SCMF. A) Integrated scRNAseq data from P7 WT, and P30 Stc1-BB het and Stc1-BB KO lung myofibroblasts. Stc1-BB KO undead SCMF (uSCMF) clusters are outlined. B) P7 WT, and P30 Stc1-BB het and Stc1-BB KO lung myofibroblasts from (A) shown by age/genotype. uSCMF clusters are outlined. C) Differential gene expression between UMAP clusters from (A). D) Comparison of DMF gene marker expression (*Cdh4*, *Igfbp5*), and development marker gene *Meox1* in P7 WT, P30 Stc1-BB Het, and P30 Stc1-BB KO DMF, SCMF, and uSCMF. Arrows for *Cdh4* and *Igfbp5* indicate expression in DMF and uSCMF. Arrows for *Meox1* indicates expression in P7 WT SCMF. E) Comparison of inflammatory marker gene expression (*Fosb*, *Icam1*, *Cxcl14*) in P7 WT, P30 Stc1-BB Het, and P30 Stc1-BB KO DMF, SCMF, and uSCMF. Arrows indicate expression in uSCMF. F) Confocal microscopy of Stc1-BB het and Stc1-BB KO lungs immunostained for SM22 at P30. SM22+ cells are from small airways. Scale bar: 20 µm.

**Supplementary Figure 6.**
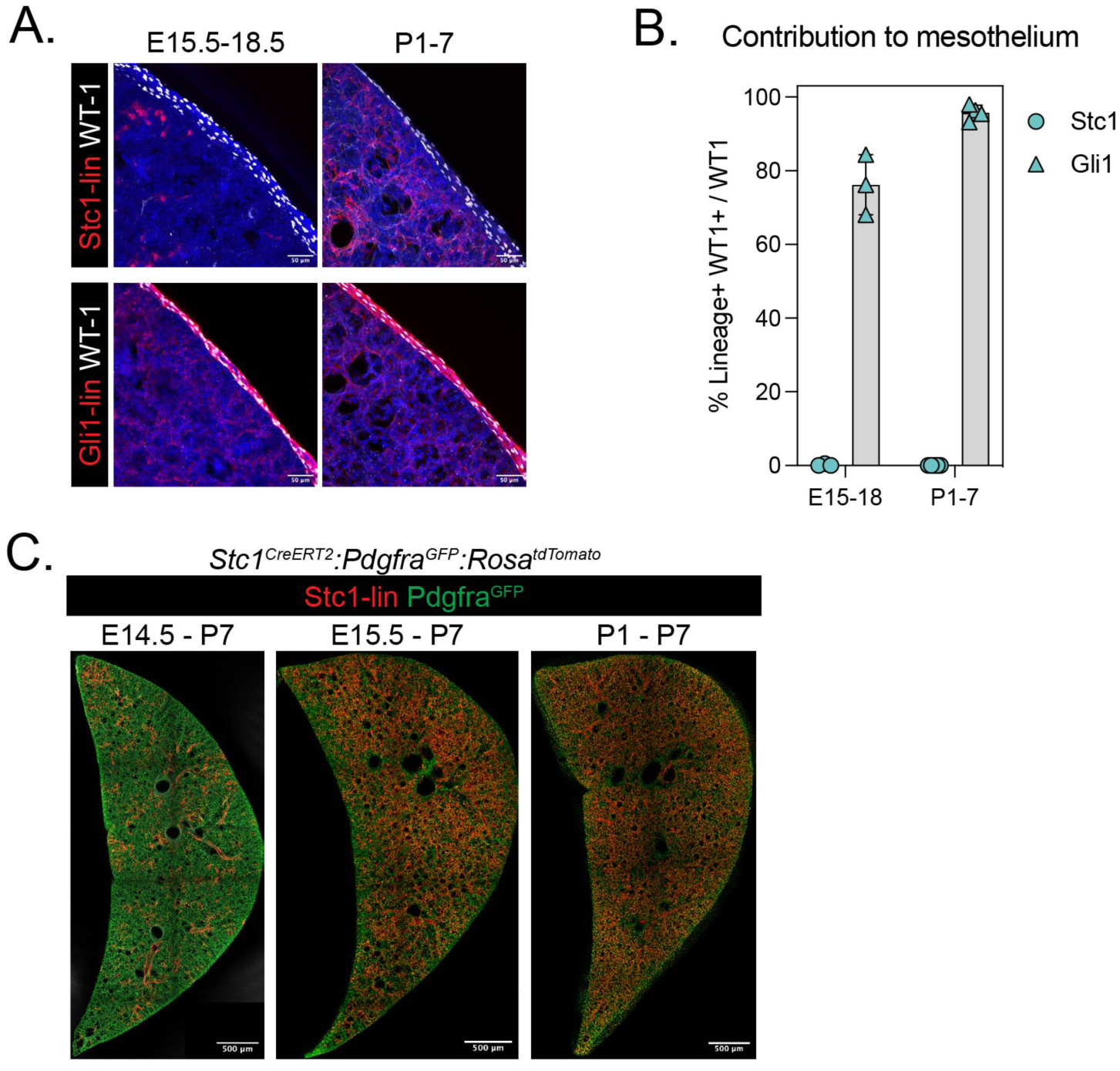
Stc1-lineage SCMF progenitors are specified by E15.5 at the RAGE-Sox9 transition zone and are distinct from Lgr6-lineage DMF and WT-1 expressing mesothelium. A) Comparison of Stc1-lineage and Gli1-lineage contributions to mesothelium immunostained for WT-1 during embryonic (E15.5-E18.5) and neonatal (P1-P7) lung development. Z-projections of approximately 100 µm. B) Quantification of (A). Data are represented as mean and standard deviation, with individual data points shown. C) Time course of Stc1-lineage labeling, analyzed for contribution to SCMF population at P7. Z-projections of approximately 80 µm. Abbreviations: Stc1-lin, Stc1-lineage; Gli1-lin, Gli1-lineage Scale bars: (A) 50 µm, (C) 500 µm.

**Supplementary Figure 7.**
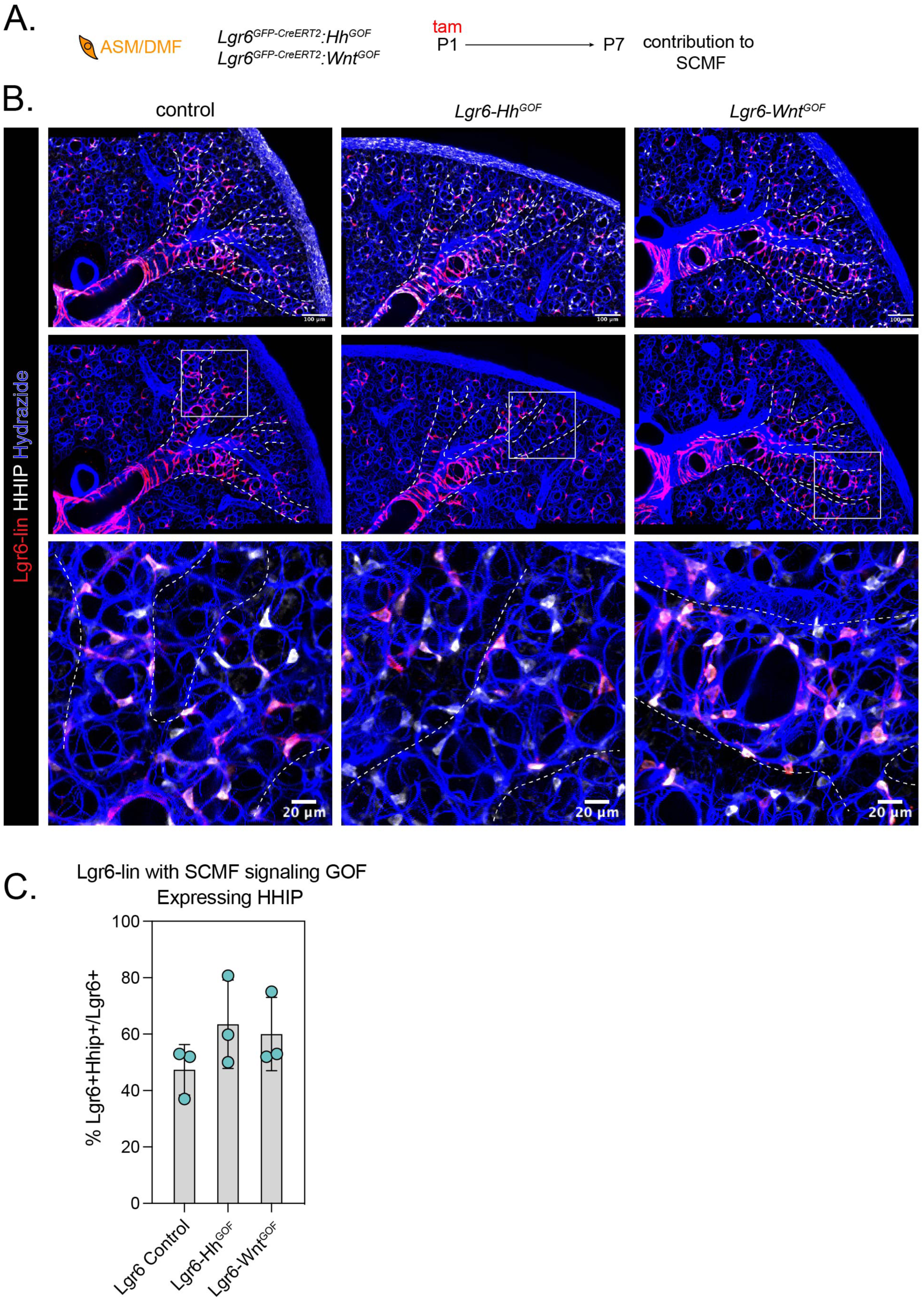
Activation of Hh or Wnt signaling in DMF progenitors does not alter cell fate. A) Diagram of genetic strategy to activate Hh and Wnt signaling pathways in Lgr6-lineage airway smooth muscle and ductal myofibroblast progenitors at P1, and analyze for any lineage conversion to SCMF at P7. B) Confocal microscopy of control, Lgr6-Hh^GOF^, and Lgr6-Wnt^GOF^ lungs at P7 immunostained for ASM/DMF marker HHIP. Z-projections of approximately 100 µm. C) Quantification from (B) of the percent of Lgr6-lineage cells that are positive for HHIP. No significant difference was found between genotypes (Kruskal-Wallis test statistic = 1.718, P value = 0.49). Data are represented as mean and standard deviation, with individual data points shown. Abbreviations: Lgr6-lin, Lgr6-lineage; Hh, Hedgehog; GOF, gain of function; tam, tamoxifen; ASM/DMF, airway smooth muscle/ductal myofibroblast; SCMF, secondary crest myofibroblast.

## SUPPLEMENTARY MOVIE LEGENDS

**Supplementary Movie 1. Stc1-lineage SCMF progenitors are specified by E15.5 at the RAGE-Sox9 transition zone.** Stc1-lineage trace from E15.5 to E16.5, immunostained with SOX9 and RAGE. Dashed lines SOX9-expressing distal tip (blue) and RAGE-expressing stalk (white) endoderm, and arrows indicate Stc1-lineage cells expressing Pdgfra^GFP^. Z-projection of approximately 65 µm.

**Supplementary Movie 2. Lgr6-lineage DMF progenitors are specified by E15.5 at the RAGE stalk. Lgr6-lineage trace from E15.5 to E16.5, immunostained with SOX9 and RAGE.** Dashed lines indicate SOX9-expressing distal tip (blue) and RAGE-expressing stalk (white) endoderm, and arrows indicate Lgr6-lineage cells. Z-projection of approximately 90 µm.

## Notes

### Competing Interest Statement

The authors have declared no competing interest.

